# A spatially-tracked single cell transcriptomics map of neuronal networks in the intrinsic cardiac nervous system

**DOI:** 10.1101/2020.07.29.227090

**Authors:** Alison Moss, Shaina Robbins, Sirisha Achanta, Lakshmi Kuttippurathu, Scott Turick, Sean Nieves, Peter Hanna, Elizabeth H. Smith, Donald B. Hoover, Jin Chen, Zixi (Jack) Cheng, Jeffrey L. Ardell, Kalyanam Shivkumar, James S. Schwaber, Rajanikanth Vadigepalli

## Abstract

We developed a spatially-tracked single neuron transcriptomics map of an intrinsic cardiac ganglion - the right atrial ganglionic plexus (RAGP) that is a critical mediator of vagal control of the sinoatrial node (SAN) activity. We developed a 3D representation of RAGP with extensive mapping of neurons and used neuronal tracing to identify the spatial distribution of the subset of neurons that project to the SAN. RNAseq of laser capture microdissected neurons revealed a distinct composition of RAGP neurons compared to CNS neuronal subtypes. High-throughput qPCR of hundreds of laser capture microdissected single neurons led to a surprising finding that cholinergic and catecholaminergic neuronal markers *Th* and *Chat* were correlated, suggesting multipotential phenotypes that can drive neuroplasticity within RAGP. Interestingly, no single gene or module was an exclusive marker of RAGP neuronal connectivity to SAN. Neuropeptide-receptor coexpression analysis revealed a combinatorial paracrine neuromodulatory network within RAGP, informing follow-on studies on the vagal control of RAGP to regulate cardiac function in health and disease.

## Introduction

In this study, we performed a spatially-tracked single cell transcriptomic analysis of an intrinsic cardiac ganglion in the pig heart to uncover the complex molecular landscape and putative paracrine neuromodulatory networks. The functional significance and complexity of the intrinsic cardiac nervous system (ICNS) has been studied for years with the majority of the focus on the physiological aspects. The ganglia at the heart are thought to constitute a “little brain” with afferent, parasympathetic and sympathetic components^1,2^. Physiological studies using epicardial ablation have demonstrated that the intrinsic cardiac ganglia mediate central control of cardiac function through vagal and sympathetic circuits^3–5^. Yet, little is known about the distribution and organization of the molecular profiles of the neurons constituting the intrinsic cardiac ganglia. Recent single neuron gene expression profiling studies have uncovered the wide range of molecularly-defined subtypes^6,7^, gradient-based organization driven by inputs to neurons^8–10^, as well as molecular plasticity during homeostatic conditions and physiological perturbations^8,9,11^ in the CNS as well as in the peripheral ganglia^7^. We set out to pursue such approaches and combine them with 3D positional information within the tissue to develop an extensive spatially-tracked molecular map of the intrinsic cardiac nervous system (ICNS).

In the present study, we focus on the right atrial ganglionic plexus (RAGP) as a key control point in the circuit that mediates vagal modulation of the SA node (SAN) activity. The routes of parasympathetic and sympathetic control of SAN were determined by several studies; surgical ablation of different areas of the heart and vagal stimulation showing a shift in pacemaker activity^12–14^. Stimulation of the RAGP fat pad was shown to cause a reduction in heart rate in dogs and human patients as well as cause a pacemaker shift^15–17^. Later studies ablating RAGP showed that this group of neurons are critical to the cardiac pacemaker response to vagal stimulation^18^. Molecular analysis of RAGP, and ICNS in general, has largely been targeted at the protein level using immunolabeling. The majority of the cardiac ganglia have been found to be cholinergic (ChAT+)^19–22^, while the proportion of observed catecholaminergic (TH+) neurons was widely variable across studies^21–23^. Additionally, some studies have reported co-expression of ChAT and TH in 10-20% of ICNS neurons^21,24^. The expression of other neurotransmitter/neuromodulator systems such as NPY, GAL, and SST have also been described within ICNS^25–30^. Here, we undertake a broad-based survey to characterize the gene expression of a wide range of neuronally relevant processes and their co-expression in single neurons with RAGP.

In a recent proof-of-principle study, we demonstrated a coordinated experimental approach that integrates imaging technologies with high throughput gene expression data (HT-qRTPCR) to develop a 3D anatomical and molecular map of rat ICNS^31^. Here, we build on that approach to incorporate single cell scale RNAseq and precisely integrate molecular data into a digitally reconstructed 3D RAGP with anatomical context of the pig heart. This approach contrasts with that of typical droplet-based single cell transcriptomics techniques in that the spatial and anatomical information of each sample is extensively tracked, which allows mapping of the molecular information into a 3D reconstructed anatomical organization of the tissue. Such an integrated anatomical-molecular map permits analysis of relationships between spatial location and molecular profiles, within the tissue as well as reference to adjacent anatomical features^9,31^ Our newly developed data demonstrate that the local cardiac ganglia harbor anatomical and molecular features necessary to function as complex signal processing units that critically mediate vagal control of heart function and health^32^.

## Results

### Mapping spatially-tracked single cell transcriptomics onto an imaging-based 3D tissue reconstruction of pig right atrial ganglionic plexus

We developed a 3D map of the single neuron scale gene expression within the pig RAGP. We used our recently developed method pipeline that combines single neuron anatomical position using 3D mapping with gene expression data of the mapped neurons obtained from single cell scale RNA-seq and High-Throughput qRT-PCR (HT-qPCR)^31^ (Fig. 1a). The RAGP neurons that project to SAN were labeled by a tracer that was injected into the SAN 3 weeks prior to sacrificing the animal for tissue harvesting. The heart tissue corresponding to the location of RAGP was sectioned serially from superior to inferior end for laser capture microdissection of single neurons and neuron pools (n=4 animals). During sectioning, blockface images were obtained, which were contoured and organized into a 3D image stack (2,698 images across n=4 animals). After sectioning and staining, tissue was subjected to laser capture microdissection where both FastBlue labeled (SAN-projecting) and unlabeled (considered as Non-SAN-projecting in the present analysis) single neurons were collected for microfluidic HT-qPCR (405 single neurons X 241 genes per neuron) and regional neuronal lifts were collected for single cell scale RNAseq (90 neuron pools). Image tracking was used to digitally annotate the spatial locations for single cell neuronal samples on a digitally reconstructed RAGP using TissueMapper software (Fig. 1b, Supplementary Movie 1). Expression levels of *Uchl1* (*PGP9.5*), a pan neuronal marker, showed a wide range that is persistent throughout the RAGP with no spatial bias for enhancement or depletion (Fig. 1b,c). We assessed all single neuron samples for expression of typical neuronal markers, cholinergic and catecholaminergic markers, and key neuropeptides (Fig. 1c-e). Nearly 100% of all collected samples not only showed detectable expression, but also abundant expression of *NeuN*, a common neuronal marker^33^. *PGP9.5* and *Map2*, another neuron-specific gene^34,35^, were also abundantly expressed in a large majority of the sampled single neurons (Fig. 1c). A high percentage of neurons showed abundant expression of *Chat, Th*, and to some extent, *Dbh*, suggesting a high degree of co-expression between cholinergic and catecholaminergic markers across single RAGP neurons (Fig. 1d). Neuropeptides such as Neuropeptide Y (*Npy*), Galanin (*Gal*), and Somatostatin (*Sst*), also showed abundant expression in a high proportion of neurons in the RAGP (Fig. 1e).

**Figure 1:**
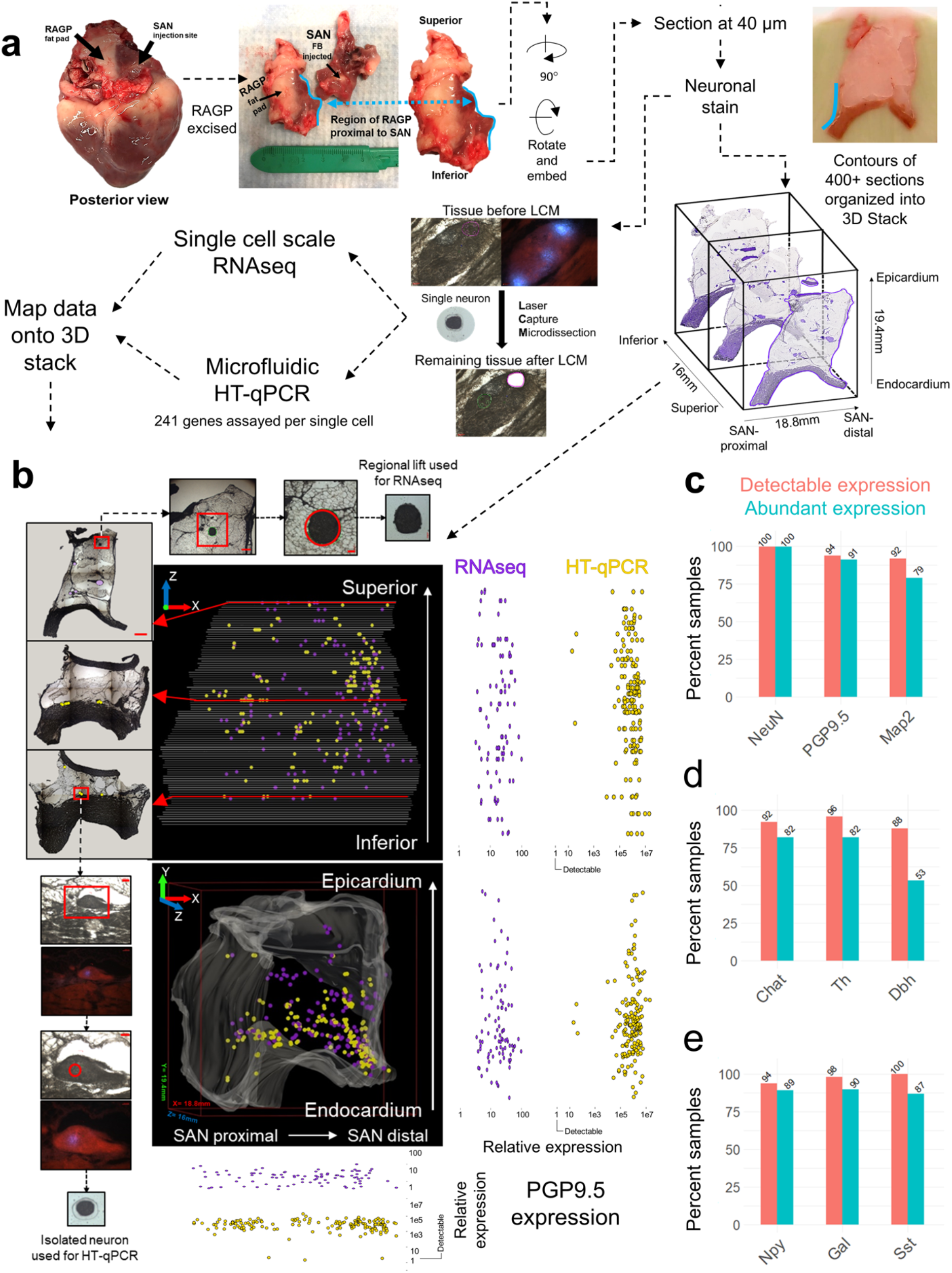
Mapping spatially-tracked single cell transcriptomics onto an imaging-based 3D tissue reconstruction of pig right atrial ganglionic plexus (RAGP). **(a)** Integrated workflow starting from injection of neuronal tracer into the sino-atrial node region of pig heart, followed by isolation, embedding, and cryosectioning of the RAGP, acquisition of block face images for 3D reconstruction, staining for neuronal localization within the tissue, and obtaining spatially-tracked single neuron samples via laser capture microdissection (LCM) for downstream processing using RNAseq and high-throughput real-time PCR (HT-qPCR), yielding transcriptomic data that is mapped onto a 3D anatomical framework. Scale bars: 50 μm **(b)** Representative visualization of the 3D anatomical framework of an RAGP depicting the location of spatially-tracked single neurons sampled via LCM (purple dots - neuronal samples for RNAseq; yellow dots - neuronal samples for HT-qPCR). The cross-sections of the stack show the corresponding tissue sections from which the neuronal samples were obtained. Scale bars: whole tissue sections, 500 μm; regional lift zoom 1, 500 μm; regional lift zoom 2, 100 μm; isolated neuron zoom 1, 100 μm; isolated neuron zoom 2, 50 μm. The relative expression of *PGP9.5* in these spatially-tracked neuronal samples is shown with reference to the axes of the 3D stack. (c,d,e) Proportion of samples that showed detectable and abundant expression of select pan-neuronal markers (c), cholinergic and catecholaminergic markers (d), and neuropeptides (e), as assessed by HT-qPCR. Data shown is based on combining 405 single neuron samples across n=4 animals.

### Transcriptomic landscape of pig RAGP from a single-cell scale RNAseq analysis

142 regional neuronal samples from the RAGP of a female Yucatan minipig were collected through LCM and spatially tracked as described above and subjected to singlecell scale RNAseq using the Smart-3SEQ protocol^36^ (described in methods). After quality check we were left with 90 samples with detectable expression in 15,000 genes. Using publicly available data in the GTEx database^37^, we retrieved a list of genes that are statistically enriched in neuronal tissues compared to other tissue types (*p* < 0.01). A total of 1,639 neuronally enriched genes showed detectable expression in our RNA-seq data (Fig. 2a). Of note, genes for ion channels associated with calcium signaling as well as glutamatergic receptors were expressed at high levels throughout the RAGP (Fig. 2b).

**Figure 2:**
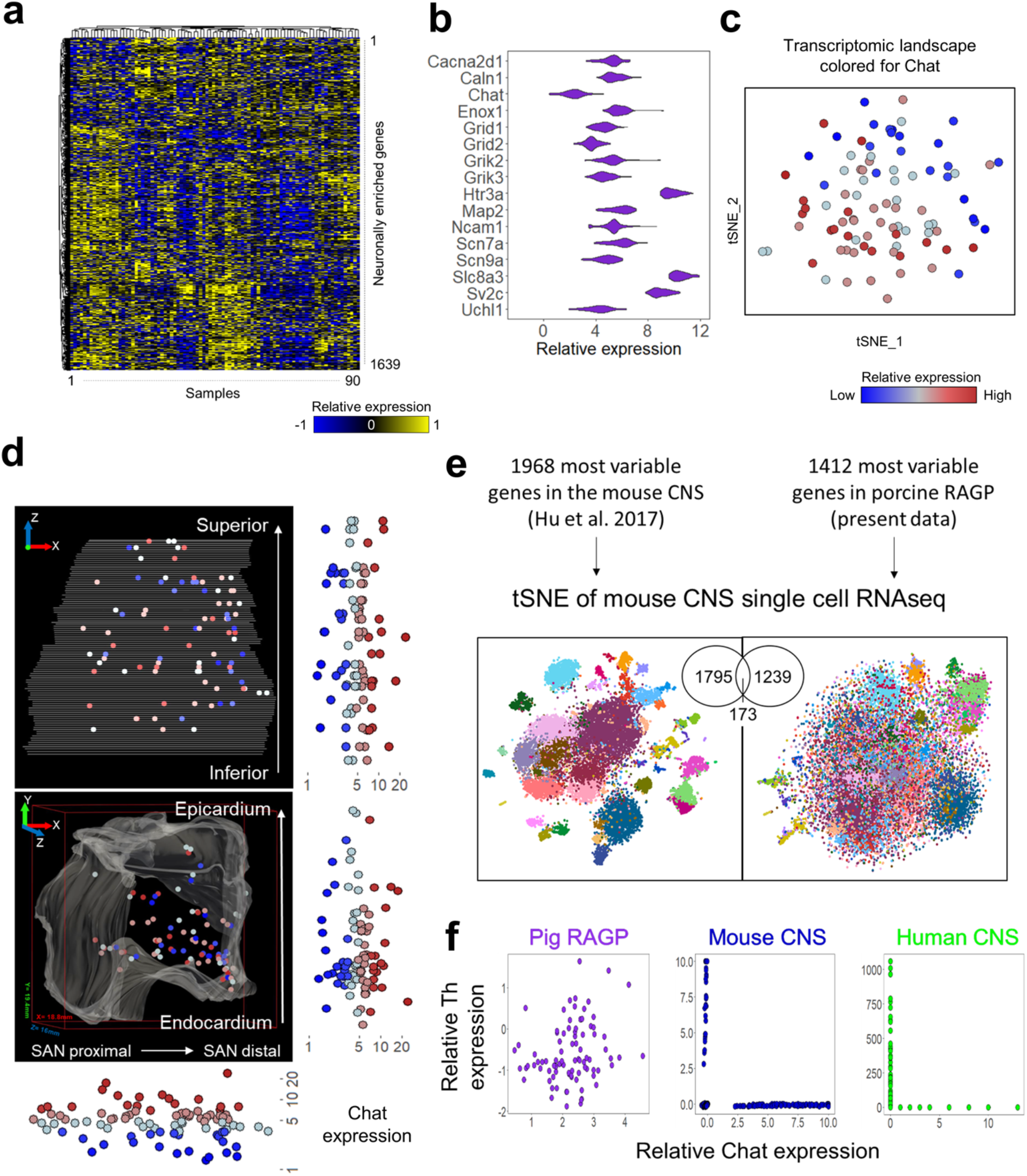
Transcriptomic landscape of pig RAGP from a single-cell scale RNAseq analysis. **(a)** Expression of 1882 neuronally-enriched genes in 90 spatially-tracked neuronal clusters in RAGP based on single-cell scale RNAseq profiling of laser capture microdissected samples. The genes included in the heatmap were selected by analyzing the GTEx database for those enriched in the neuronal tissues compared to other tissue types. Of the genes identified as neuronally enriched in the GTEx database, 1639 genes were present in the RAGP neurons. **(b)** Distribution of select abundantly-expressed neuronal genes. **(c)** Transcriptomic landscape as delineated by tSNE indicating the gradient of *Chat* throughout a distributed cloud. **(d)** Visualization within the 3D anatomical framework for a representative RAGP. The relative expression of choline acetyltransferase (*Chat*) in these spatially-tracked neuronal samples is shown with reference to the axes of the 3D stack. **(e)** A comparison of the distribution of CNS neuronal types based on the most-variable genes in mouse CNS (Hu et al., 2017) versus in pig RAGP (present data). The tSNE plots are colored based on 40 distinguishable mouse CNS neuronal states described in Hu et al. (2017). **(f)** Scatter plots comparing the expression of *Th* vs *Chat* in the pig RAGP (present data), mouse CNS^6^, and human CNS^39^ (https://portal.brain-map.org/atlases-and-data/rnaseq/human-multiple-cortical-areas-smart-seq).

The transcriptomic landscape of the neuronally enriched sample set was assessed through a nonlinear embedding approach, t-stochastic neighbor embedding (tSNE)^38^. The results show that the samples were not separated into groups corresponding to distinguishable phenotypes, but were instead organized as a single cloud suggesting a gradient of gene expression based organization of underlying neuronal molecular states^8^. Coloring the tSNE map for relative expression of *Chat*, an important marker for cholinergic expression, reveals a distinct gradient across the transcriptomic landscape (Fig. 2c). We then visualized the expression pattern of *Chat* in the context of 3D anatomical space. Similar to the spatially distributed expression of *PGP9.5* (Fig. 1b), *Chat* expression was widely distributed throughout the RAGP with no spatial gradients along any axis (Fig. 2d).

We compared the transcriptomic profiles of the RAGP neurons with the molecular phenotypes identified from single neuron transcriptomics analysis in the CNS^6^. We used the available mouse CNS data, due to lack of comparable pig CNS single neuron transcriptomic data set. We reconstructed the tSNE map of single neurons in the mouse CNS based on the 1,968 expression of the highly variable genes^6^ (Fig. 2e). To compare how the neuronal phenotypes identified in the CNS compared to those in the RAGP, we first extracted genes contributing most to the variability in the RAGP through Principal Component Analysis (PCA). Of the 1,814 most variable genes in the RAGP, a subset of 1,412 genes were found in the mouse CNS single neuron data^6^. Visualization by tSNE map shows that the well defined neuronal clusters based on variable genes in the CNS are largely lost when the neuronal heterogeneity was analyzed using the 1,412 most variable genes in the RAGP (Fig. 2e). These results demonstrate that the highly variable genes within RAGP may not be as variable in the CNS and vice versa, suggesting a different organization of neuronal heterogeneity between RAGP and CNS structures.

Considering that a large proportion of neurons showed expression of *Th* and *Chat* (Fig. 1d), we examined the co-expression patterns of these genes in the transcriptomics data. Neurons in the mouse^6^ and human CNS^39^ show mutually exclusive expression of *Th* and *Chat*. In stark contrast to the lack of *Th* and *Chat* co-expression in the CNS, we found a high degree of co-expression between *Th* and *Chat* in the RAGP neurons (Fig. 2f). This finding further augments the results seen in the tSNE map (Fig. 2e) suggesting that the neuronal molecular states within the RAGP may be organized in a manner that is not similar to the neuronal phenotypes observed in the CNS.

### Landscape of neuronal transcriptional states in the pig RAGP

Using the 3D map of a representative RAGP, examination at the single cell scale allowed us to visualize the distribution of neurons within their 3D anatomical framework, revealing that neurons, while distributed throughout the RAGP, are more densely packed closer to the endocardium (Fig. 3a). Annotation of neurons based on their projection to the SAN indicates that while both projecting and non-projecting neurons are present throughout the RAGP, SAN-projecting neurons appear to be more concentrated closer to the SAN and less concentrated towards the epicardium (Fig. 3a-c, Supplementary Movie 1). We assayed 405 spatially-tracked single neuron samples from RAGP (n=4 animals) for expression of 211 genes within each neuron using HT-qPCR, representing both SAN-projecting and non SAN-projecting neurons (Supplementary Fig. 1a). At the molecular level, a set of only 6 genes (*Cck, Gal, Grp, Hcrtr1, Ntrk1, Ret*) showed significant differences in the distribution of expression between SAN-projecting and SAN non-projecting neurons (K-S statistic, FDR-adjusted *p* < 0.01, fold change > 2, Supplementary Fig. 1b). Comparing neurons across all 4 RAGP, the expression distribution of select neuronal markers *NeuN, PGP9.5, Chat, Th, Dbh*, and *Npy* were relatively consistent across animals and between SAN-projecting and non SAN-projecting neurons (Supplementary Fig. 1c).

**Figure 3:**
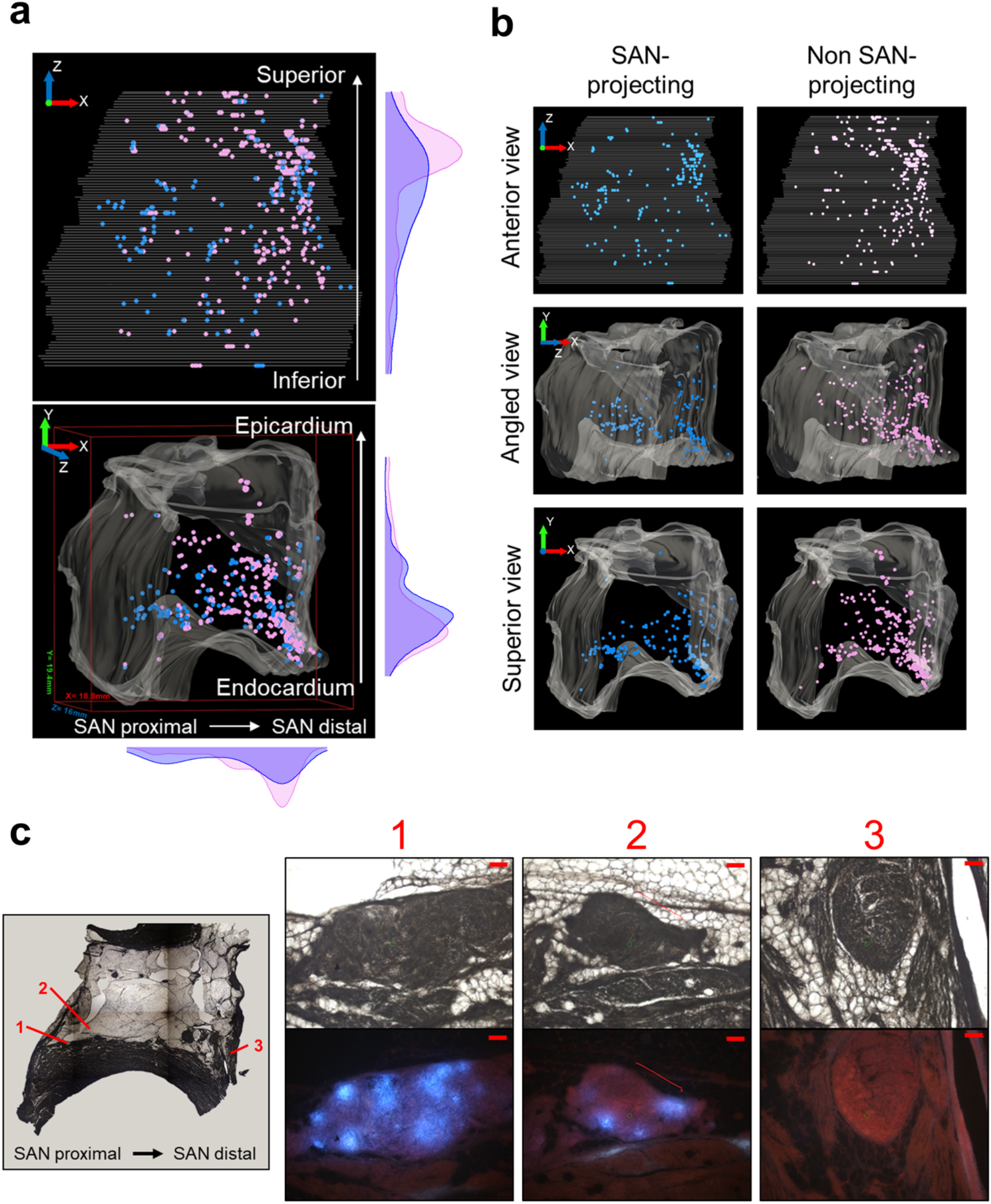
Broad RAGP anatomy of SAN-projecting and non SAN-projecting neurons. **(a)** Visualization within the 3D anatomical framework of both SAN-projecting (blue) and non SAN-projecting (purple) neurons that were comprehensively identified in select sections of a representative RAGP. Panels along the right side and bottom show density plots representing the density of projecting and non-projecting neurons along each axis. **(b)** Anterior (top), angled (middle) and superior (bottom) views of a representative RAGP showing only the SAN-projecting neurons (left) or non SAN-projecting neurons (right). **(c)** A select section of the RAGP (7,040 μm from the superior aspect) zooming in on three different neuron clusters showing a high percentage of neurons within the cluster projecting to the SAN towards the SAN-proximal side of the RAGP (1), a cluster with no SAN-projecting neurons towards the SAN-distal side of the RAGP (3) and a cluster with a mix of both projecting and non-projecting neurons in between (2). Scale bars: 100 μm. For an animated visualization see Supplementary Movie 1.

In order to identify gene expression modules that can better characterize the neurons based on the connectivity to SAN, we used a combination of clustering and template matching. Our approach yielded six transcriptional states within the RAGP neurons (Fig. 4a). Briefly, the gene expression profiles of SAN-projecting neurons from one male and one female RAGP were subjected to hierarchical clustering to partition the single neurons into distinct states. These states were used as templates to assign the remaining non SAN-projecting neurons to one of these states based on correlation to the template, with neurons below the correlation threshold sequestered into an additional state. Neurons from a different pair of male and female RAGP were sorted into these transcriptional states based on correlation to the template profile, revealing similar patterns (Fig. 4a, Supplementary Fig. 2). Examination of these transcriptional states reveals that the molecular signatures of SAN-projecting and non-SAN projecting neurons are remarkably similar. A closer examination uncovered a state consisting of almost entirely SAN-projecting neurons (state C), and another state consisting of almost entirely SAN non-projecting RAGP neurons (state F). Notably, neither of these states could be distinguished from the other neuronal states by the patterns of any given gene expression module. Instead, each state was characterized by a combinatorial pattern of multiple gene expression modules (Fig. 4a). Visualization using a tSNE map revealed that within the context of the transcriptional landscape, these neuronal states are distributed across parts of a single cloud, suggesting a gradient-based organization of the neuronal states (Fig. 4b). Analysis of the potential patterns and gradients of these states within the 3D anatomical framework of a representative RAGP revealed that the neuronal states were evenly distributed throughout the RAGP (Fig. 4c, Supplementary Movie 2).

**Figure 4:**
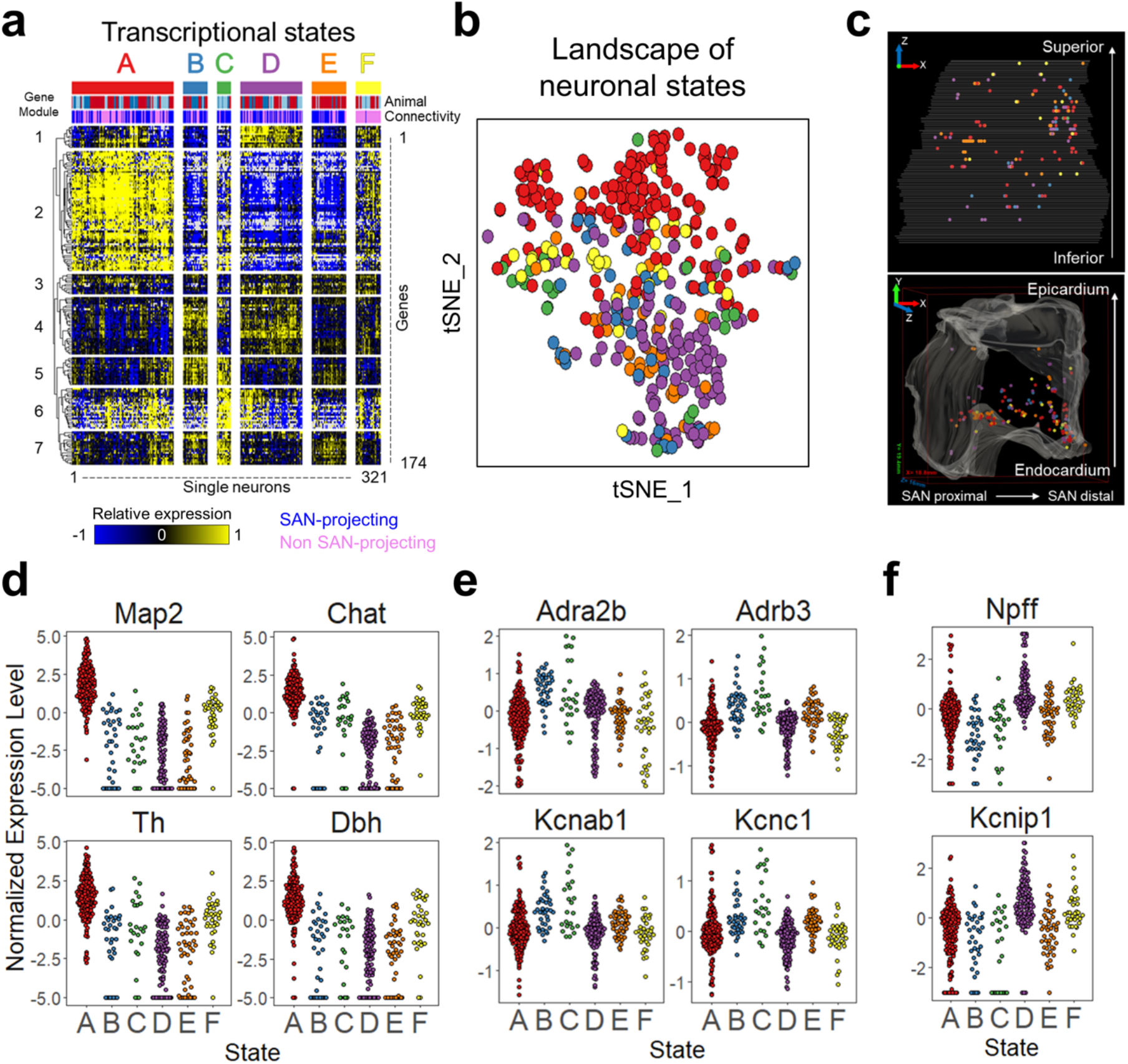
Landscape of neuronal transcriptional states in the pig RAGP. **(a)** Expression of 174 genes, each assayed in 321 single neurons (n=3 animals), yielding six transcriptional states using a combination of clustering and template matching analysis. A majority of the states consisted of both SAN-projecting and non SAN-projecting neurons. A complete heatmap with 405 neurons from all 4 RAGP is shown in Supplementary Fig. 2. **(b)** Landscape of neuronal transcriptional states visualized as a tSNE plot. Colors correspond to the states shown in panel (a). **(c)** Visualization of the neuronal states within the 3D anatomical framework for a representative RAGP. **(d-f)** Expression distribution of select genes with enrichment in specific transcriptional states: (d) state A - *Map2, Chat, Th, Dbh*; (e) states B and C - *Adrab2b, Adrb3, Kcnab1, Kcnc1*; and (f) states D and F - *Npff* and *Kcnip1*. Enrichment assessed by a one-way ANOVA and post hoc Tukey Honest Significant Difference test p-value < 0.01.

We further explored the heterogeneity of gene expression distribution across these neuronal states (Fig. 4d-f, Supplementary Fig. 1a, 3a). State A is largely defined by high expression of genes in module 2 which include neuron-specific genes such as *Map2, Eno2*, and *PGP9.5* as well as genes involved in neurotransmitter processes such as *Chat, Th*, and *Dbh* (Fig. 4d). States B and C were both characterized by high expression of genes in modules 5 and 7, which include several adrenergic receptors and potassium channels (Fig. 4e). Modules 4 and 6 delineate the separation between states B and C where module 4 is characterized by high expression in states B and D and includes a variety of receptor subtypes. Module 6, consisting of a variety of neuropeptides and neuropeptide receptors, has a broader range of expression across all samples, with mixed expression in most states but particularly high expression in state C. Genes in module 1 were upregulated in states D and F and included neuropeptides such as *Npff* and *Nppa* as well as a voltage-gated potassium channel, *Kcnip1* (Fig. 4f). Examination of ion channels and glutamatergic and GABAergic receptors also show a range of expression across the identified transcriptional states (Supplementary Fig. 3b,c).

### Correlated cholinergic and catecholaminergic gene expression in RAGP neurons

Examination of genes involved in acetylcholine and catecholamine biosynthesis and transport processes shows consistent expression across all RAGP and between SAN-projecting and non SAN-projecting neurons and also reveal a surprising level of coexpression (Fig. 5a,b, Supplementary Fig. 1a, 4a-d). A closer look at the genes involved in the catecholamine biosynthesis showed that *Th*, the rate limiting enzyme in the production of all catecholamines, and *Dbh*, responsible for the conversion of dopamine to norepinephrine are expressed over a wide range and show a high degree of coexpression (Figure 5b, Supplementary Fig. 4b). *Ddc*, the enzyme converting L-DOPA to dopamine, and *Pnmt*, responsible for the conversion of norepinephrine to epinephrine, were both stably expressed in the majority of samples as seen by the high abundance and narrow range of expression for both genes (Fig. 5b). This further underscores the regulation of catecholamine biosynthesis process at the level of *Th* and *Dbh* as the rate-limiting enzymatic steps whose gene expression levels are highly variable across single neurons.

**Figure 5:**
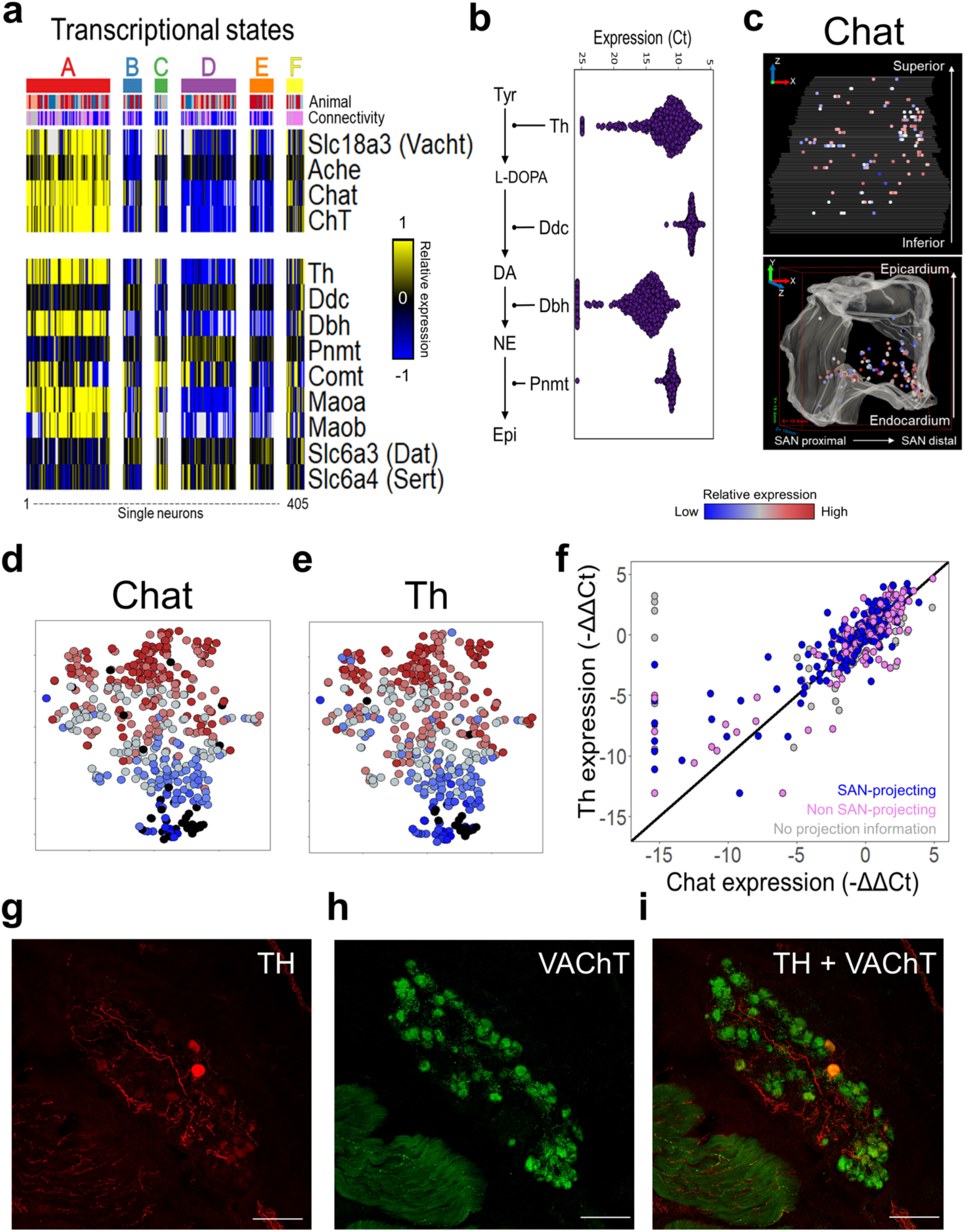
Correlated cholinergic and catecholaminergic gene expression in RAGP neurons. **(a)** Transcriptional state-wise gene expression of the components of acetylcholine and catecholamine biosynthesis and transport processes, across 405 single neurons in RAGP. **(b)** Beeswarm plot showing the abundance and the range of expression of key genes involved in catecholamine biosynthesis across 405 single RAGP neurons (n=4 animals). **(c)** Visualization of *Chat* gene expression within the 3D anatomical framework for a representative RAGP. **(d,e)** The distributions of *Chat* and *Th* gene expression overlap within the transcriptional landscape as visualized in the tSNE plots. **(f)** Correlated gene expression of *Chat* and *Th* across single neurons in RAGP (R^2^=0.69, p-value < 2.2e-16). The pair-wise comparison of gene expression levels is shown for SAN-projecting and non SAN-projecting neurons (n=4 animals). The points marked grey correspond to RAGP neurons without information on SAN projection, as these were microdissected from a pig heart without a tracer injection into the SAN region. **(g,h,i)** Confocal images showing a cluster of neurons within RAGP double stained for TH (g) and VAChT (h). Colocalization of TH and VAChT in a subset of neurons (i).

Visualization of key genes such as *Chat* and *Th* within the 3D anatomical framework of a representative RAGP revealed that their wide range of expression is distributed spatially throughout the RAGP (Fig. 5c; Supplementary Figure 5a, Supplementary Movie 3). Visualization of the expression distribution of *Chat* and *Th* through a tSNE map revealed distinct and overlapping gradients within the transcriptional landscape (Fig. 5d,e). Single cell scale RNAseq data suggested regional co-expression of Th and Chat (Fig. 2f). Building on these results, single neuron gene expression analysis showed that *Chat* and *Th* were highly correlated across single neurons in the RAGP (Fig. 5f). Interestingly, gene co-expression of these cholinergic and catecholaminergic markers was in stark contrast to protein expression patterns that showed much reduced overlap of expression between TH and VAChT, another cholinergic marker, in individual neurons (Supplementary Fig. 4c). Immunohistochemistry for TH and VAChT revealed that a majority of neurons showed robust protein expression of VAChT with a subset co-staining for TH (Fig. 5g,h,i). This finding suggests that post-transcriptional regulation plays a key role in shaping the neurotransmitter patterns within RAGP. In particular, the high correlation between *Th* and *Chat* at the mRNA level suggests that the RAGP neurons are poised to use both cholinergic and catecholaminergic processes, along the lines of multipotential neuronal phenotypes observed in CNS^8^.

### Neuropeptidergic interaction networks in the pig RAGP

We examined the co-expression patterns of neuropeptides and their cognate receptors to identify putative local paracrine networks within RAGP. Identifying neuronal subsets based on co-expression patterns of a neuropeptide with its receptors allows us to classify groups of neurons that exhibit autocrine or paracrine signaling. We examined local paracrine networks for three important neuropeptides *Sst, Gal*, and *Npy* in detail (Fig. 6a-f, Supplementary Fig. 5b-d,6).

**Figure 6:**
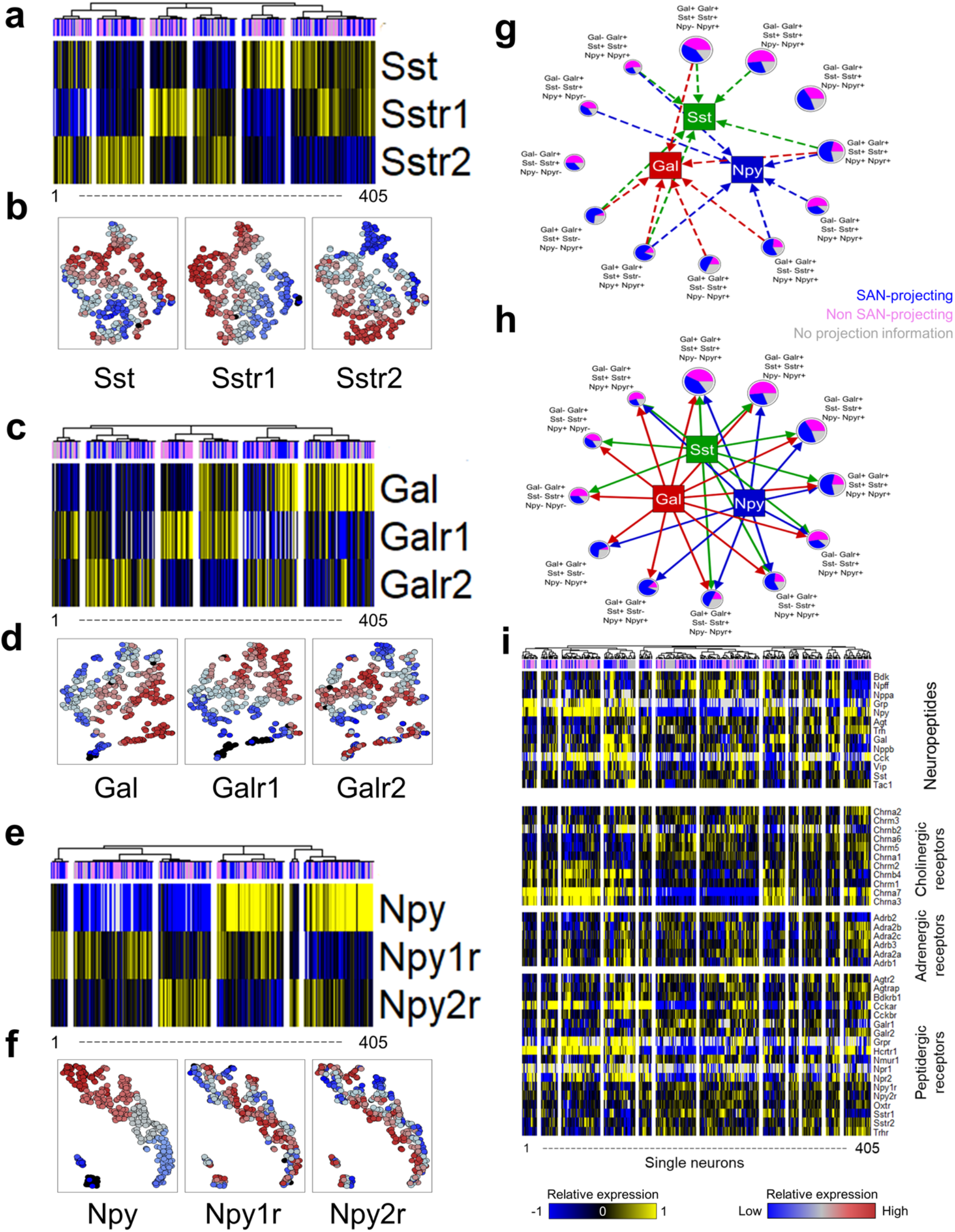
Neuropeptidergic interaction networks in the pig RAGP. **(a,c,e)** Expression patterns of somatostatin (a), galanin (c), and neuropeptide Y (e) and their cognate receptors across 405 single RAGP neurons (n=4 animals). **(b,d,f)** The transcriptional landscape of the neurons varies based on the selected neuropetide-receptor set. The distribution of expression of the neuropeptides and their receptors shown in the tSNE plots (b,d,f) is enriched in distinguishable parts of the landscape, corresponding to the combinatorial expression seen in the heat maps (a,c,e). **(g,h)** Interaction networks of neuronal subtypes defined based on the combinatorial pattern of neuropeptides and their receptor expression. **(g)** The interaction network subset corresponding to the neuronal subtypes producing the neuropeptides Somatostatin (*Sst*), galanin (*Gal*), and neuropeptide Y (*Npy*). The circular nodes denote the neuronal subtypes.The size of the node is proportional to the number of single neurons belonging to each subtype. The pie chart within each circular node indicates the proportion of the neurons within that subtype that are identified as projecting to the SAN region. The arrows from the circular nodes denoting the neuronal subtypes connect to the square-shaped nodes denoting the three neuropeptides, based on which subtypes show the corresponding neuropeptide gene expression above a specified threshold. The color of the arrows matches the color of the corresponding target square-shaped node. **(h)** The interaction network subset corresponding to the neuropeptide receptor expression across the neuronal subtypes. The notation of circular and square-shaped nodes is the same as in pane (g). The arrows connect each neuropeptide to the neuronal subtypes based on which subtypes express any the corresponding neuropeptide receptors above a specified threshold. The color of the arrows corresponds to the color of the neuropeptide node. **(i)** Combinatorial pattern of expression of a wide range of neuropeptides and receptors of neurotransmitters across single neurons in RAGP (n=4 animals).

In the case of somatostatin and its receptors, *Sstr1* and *Sstr2*, neurons cluster into six groups based on the presence or absence of each gene (Fig. 6a). This revealed one neuronal subset that expresses somatostatin, but not its receptors, and therefore can transmit but not respond to the somatostatin signal in the RAGP network. Meanwhile, two neuronal subsets were positive for somatostatin as well as either receptor *Sstr1* or *Sstr2*, representing groups that can both transmit and be activated by somatostatin.The remaining three neuronal subsets do not synthesize but can respond to somatostatin (Supplementary Fig. 6a,g). Similar to *Sst, Gal* and *Npy* and their cognate receptors each clustered into six neuronal subsets. When categorizing each neuronal subset based on the presence or absence of each gene, however, the six neuronal subsets were reduced to 5 categories for the *Gal/Galr1/Galr2* set and 4 identifiers for the *Npy/Npy1r/Npy2r* set (Figure 6c,e, Supplementary Fig. 6c,e,h,i).

Examining the tSNE transcriptomic landscape filtered for the selected neuropeptide-receptor set showed distinct and overlapping patterns of expression for the genes in each set (Fig 6b,d,f). This further underscored the distinctive combinatorial expression patterns seen within each neuropeptide-receptor network. Despite the strong patterns of expression seen within each individual network, none of the genes in either the *Sst/Sstr1/Sstr2* or *Gal/Galr1/Galr2* set showed a discernible gradient across the broader transcriptomic landscape (Supplementary Fig. 6b,d,f). However, *Npy* (and not its receptors) showed a gradient of expression that largely mirrored the *Th* and *Chat* gradients, consistent with the co-expression of these genes across single neurons (Supplementary Fig. 6f). Taken together, the results indicate that while *Sst, Gal*, and *Npy* each have distinct co-expression patterns that outline local paracrine networks, the signalling of any one neuropeptide alone is unlikely to be the main driver of transcriptional states of RAGP neurons.

Integrating all three paracrine networks at once revealed that the overall combination is more complex than any individual neuropeptide driven network (Fig. 6g,h, Supplementary Fig. 6). Each neuron was categorized as neuropeptide +ve or neuropeptide -ve, receptor +ve or receptor -ve, based on whether the gene expression of the given peptide/receptor was above the median expression value for that gene across all the neurons. Applying such a categorization to the 405 laser capture microdissected RAGP neurons resulted in 61 distinct neuronal categories (out of a possible set of 64), indicating a combinatorial network of neuropeptide signalling (Supplementary Cytoscape Network File). Of these 61 categorized neuronal states, a set of 12 states consisted of more than 10 sampled neurons each, representing a total of 224 out of the 405 single neurons (about 55%). A network representation of these 12 neuronal subsets highlighted the putative autocrine and paracrine network of neuropeptide signalling between RAGP neurons (Fig. 6g,h). Visualization of these individual networks within their 3D anatomical framework also suggests the potential for widespread signaling throughout the RAGP (Supplementary Fig 5b-e, Supplementary Movie 4). Further expanding the network analysis to include all assayed neuropeptides and receptors revealed a combinatorial mosaic of neuropeptide/receptor expression (Fig. 6i), indicating a complex local neuromodulatory network within RAGP.

## Discussion

In this study, we performed a combination of spatially-tracked single cell transcriptomic analysis of an intrinsic cardiac ganglion. Our results uncovered the complex molecular landscape and paracrine networks of RAGP neurons in the pig heart. RNAseq analysis revealed that the composition of RAGP neurons is distinct from that of neuronal subtypes in the CNS, with minimal overlap in the combinatorial expression patterns of neuronally enriched genes. Using HT-qPCR of hundreds of single RAGP neurons, we identified neuronal transcriptional states with distinct gradients across the transcriptomic landscape. A remarkable finding was a high prevalence of coexpression of cholinergic and catecholaminergic neuronal markers *Chat* and *Th* in a large fraction of the RAGP neurons, suggesting multipotential phenotypes. The gene expression profiles of SAN-projecting RAGP neurons were distributed across multiple neuronal states without a single gene or gene expression module serving as an exclusive marker of RAGP neuronal connectivity to the SAN region. Our integrative analysis revealed complex expression patterns of neuropeptide signaling such that any individual neuropeptidergic system is unlikely to act as the main driver of neuronal transcriptional states. The single neuron transcriptomic landscape and paracrine networks uncovered in this study form the foundation for investigating the neuromodulatory processes that shape the RAGP neuronal network response to vagal inputs to regulate cardiac function in health and disease.

In recent years, the single cell RNASeq studies of neurons isolated from various CNS components show distinct separations along the transcriptomic landscapes with individual clusters specifically linked to excitatory or inhibitory processes^6,7,40–42^. Given the complex neural networks to and from the CNS that influence the ICNS, we compared our RNASeq data with RNASeq data from single CNS neurons to compare cell types across central and peripheral neurons. Interestingly, our data showed little to no alignment with the cell populations identified in CNS neurons. In contrast with CNS neurons which cluster distinctly into dedicated phenotypes, neurons of the RAGP resemble mixed neuronal types with a gradient of expression as represented by a single cloud across the transcriptomic landscape (Figure 2c, 4b)^6,7^. Transcriptomics and proteomics with 3D anatomical location tracking have been established for the brain (Allen Brain Atlas)^10,43–45^.To our knowledge, we are the first group to attempt this at the mammalian heart^31^. This work enabled creation of a 3D model of pig RAGP that precisely integrates molecular data of single neurons into the 3D anatomical framework. Unlike the CNS, where molecular profiles correspond to specific anatomically located nuclei, we have found no discernable connection between the molecular profiles and anatomical locations within the RAGP^10,43–45^. This suggests that the physiological functions attributed to the ICNS are likely to not be restricted to a particular anatomical location within any ganglionic plexus, and that the RAGP, and possibly other ganglionic plexuses in the ICNS, is composed of a combination of neuronal types whose integrative control of the heart enables the complex response patterns observed in physiological studies^46^.

Available physiological data suggest the notion that RAGP may consist of 80% locally connecting neurons, with the remainder distributed between those receiving afferent and motor input (both parasympathetic and sympathetic)^2,47–49^. Our results showed that the underlying transcriptomically-defined molecular states of neurons in RAGP are not necessarily aligned with the phenotypic categories postulated by physiological findings. We identified six transcriptional states of RAGP neurons that are distinguished by combinatorial patterns of gene expression modules that span several neuronal functions/phenotype categories, including cholinergic, adrenergic, neurotransmitter, receptor expressions and ion channels. No state was exclusively represented by individual gene expression modules or a subset of the genes. Such combinatorial patterning of neuronal states was present similarly in SAN-projecting and SAN non-projecting neurons. Analysis of 3D spatial location of connectivity-based (SAN projecting, SAN non-projecting) and transcriptional state-based neuronal groups showed a heterogeneous distribution across the anatomical and transcriptional landscape of the RAGP. Our results on the combinatorially organized gene expression modules defining the spatially distributed RAGP neuronal states can serve to explain the variability of physiological responses observed in experimental disruption of ICNS circuits^3–5^.

Distinct neuron types that are sympathetic or parasympathetic have been identified in the ICNS in various immuno-histochemical studies for ChAT and TH to identify cholinergic or adrenergic phenotypes, respectively^19,21–23^ where the majority have found the neurons to be either parasympathetic or sympathetic in nature. However, some have found that ICNS neurons are not exclusively parasympathetic or sympathetic, with reports showing 10-20% of the cardiac GP co-expressing both ChAT and TH^21,24^. One of the more significant findings of this study is the co-expression of cholinergic and catecholaminergic pathway genes in our data (Fig. 5). To our knowledge, this is the first study to report co-expression of both *Chat* and *Th* in the RAGP, with 90-95% of neurons showing detectable expression of both genes, and 76% of neurons showing abundant expression of both genes at the single-cell transcriptional level (Fig. 5 a-h). Of note, the ICNS is comprised of two neuronal types, large principle neurons (PNs) and small intensely fluorescent (sif) cells, the later of which contain a wide range of neurotransmitters and are often more likely to express adrenergic phenotypes^1,21,50^. Based on neuron size and through the use of LCM, we specifically targeted our single cell sampling to PNs (Fig. 3c). While it is possible that some sif cells were collected along with the PNs, we believe that the data and therefore the coexpression patterns between *Chat* and *Th* can be attributed mainly to the PNs in the RAGP. Testing the protein level expression of these enzymes in the RAGP showed consistent results with previous studies in the ICNS. Confocal microscopy studies of RAGP showed that 99% neurons were VAChT positive and 10-20% neurons also positive for TH (Fig. 5 g-i). We interpret these differences between gene and protein expression levels as representative of multipotential phenotypes that are observed at the transcriptional scale that then are shaped further post-transcriptionally to yield a specific distribution in a given physiological context. We and others have shown that cells retain their plasticity beyond development and are able to adapt to perturbation based on the inputs they receive in normal and diseased states by transcriptional, as well as post-transcriptional regulation^8,11^. For example, we have previously shown that TH immunoreactive neurons in the brainstem are organized along a gradient of catecholaminergic (*Th+/Fos-*) and non-catecholaminergic (*Th-/Fos+*) neuronal states^8^, and this gradient shifts in response to physiological perturbation such as sustained hypertension. Our results on correlation of *Th* and *Chat* raise an intriguing possibility of such plastic adaptive dynamics within the RAGP, with functional effects on vagal control of cardiac function.

We delved into three peptidergic systems, *Gal, Sst*, and *Npy*, which displayed distinctive coexpression patterns with their respective receptors. These neuromodulators have been shown to have important effects on cardiac function and vagal tone^26,27,29,51–54^. High concentrations of *Sst* have been found in cardiac tissues, specifically in the right atrium and the atrioventricular node, where specialized conducting and pacemaker cells are found^30^. *Sst* is found mostly in GABAergic neurons and mediates antagonism of sympathetic processes on a broad scale as well as promotes parasympathetic effects, specifically reducing cardiac contractility in an Ach-dependent manner^54^. Meanwhile both *Gal* and *Npy* have been identified as co-transmitters in adrenergic neurons and have been shown to work in similar and complementary manners with respect to vagal control^26^. While *Gal* has been shown to attenuate cardiac vagal activity with no effect on blood pressure, *Npy* does just the opposite, increasing blood pressure but having no inhibitory effects on vagal activity^51^. Studies have shown that *Gal* and *Npy* released from sympathetic neurons inhibit the release of acetylcholine in the cardiac cholinergic postsynaptic neurons^28^. Our results on the expression of *Sst*, *Gal*, and *Npy* in the RAGP neurons suggest a strong likelihood of these neuromodulatory interactions arising from within the RAGP paracrine networks to mediate parasympathetic and sympathetic control of cardiac function. This data is largely consistent with our previous findings in the rat heart that also show a wide range of expression and combinatorial patterns between peptides and their cognate receptors^31^. We examined each of these peptidergic networks separately as well as in combination with one another in the larger context of the neuronal network. While examination of one neuropeptide-receptor set at a time reveals discrete and overlapping co-expression patterns, these gradients are absent from the broader transcriptomic landscape, further underscoring that examination of one peptidergic network is insufficient to gain an understanding of the network as a whole. Combination of the three peptidergic networks yielded a wide range of combinatorial gene expression patterns and reveals a widespread signalling network where almost all sampled neurons are capable of being activated by one or more neuropeptides. *Gal* and *Npy* have been shown to be released together in sympathetic neurons of the stellate ganglia^28^. We found that these two neuropeptides showed partly overlapping gene expression patterns across RAGP neurons, suggesting distinctive adaptive and neuromodulatory phenotypes in the RAGP versus elsewhere.

Expanding the network analysis to account for the wide range of neuropeptide systems expressed within RAGP suggests a conceptual formulation of RAGP as a highly adaptive and dynamic system driven by combinatorial patterns of neuromodulators acting in local paracrine networks. Future studies can inform interventions that rely on the combinatorial dynamics of these paracrine networks to influence central regulation of cardiac function. Our data demonstrates that the local cardiac ganglia harbor anatomical and molecular features necessary to function as complex signal processing units that critically mediate vagal control of heart function and health. Integrating these findings with other data obtained from our collaborative SPARC consortium will enable development of computational models of neurons and networks in the RAGP as a critical mediator of vagal control of the heart^32^. With the RAGP as a starting point, the RAGP network model can be further expanded into a systems model of autonomic control of cardiovascular function to test novel control strategies for neuromodulation and open the door to more effective therapeutics.

## Methods

### Animals

Animal experiments were performed in accordance with the UCLA Institutional Animal Care and Use Committee, and euthanasia protocols conform to the National Institutes of Health’s Guide for the Care and Use of Laboratory Animals (2011). Data for all experiments were collected from male and female Yucatán minipigs (> 3 months old).

### Optimization of neural tracers for laser capture microdissection

Fast Blue (Polysciences, 17740-1), CM-DiI (Thermo Fisher, C7000), Fast DiI (Thermo Fisher, D3899), TMR-Dextran (Thermo Fisher, D3308), FluoroGold (Fluorochrome) as candidate fluorescent tracers were scanned for their compatibility with the dehydration protocol of laser capture microdissection before the tracer was adapted in the minipig. In each animal, one of the five tracers was injected into Sprague Dawley rats (3-4 months old, purchased from Envigo) sino-atrial node at the same volume of 10 μL and the heart tissues were harvested in the time window 10 am-12 pm 14 days after injection and stored in OCT immediately. Cryosections were visualized under a fluorescence microscope before and after the dehydration steps necessary for laser capture microdissection and acquisition of single neuron samples (as detailed below). All five tracers labeled intrinsic cardiac nervous system neurons successfully but only CM-DiI and Fast Blue-labeled neurons remained intact on the heart sections without fixation, whereas TMR-Dextran and FluoroGold-stained neurons were not visible under the microscope under the conditions suitable for laser capture microdissection. Only Fast Blue provided reliable and consistent labeling visible under laser capture microdissection microscope after the necessary dehydration procedure and was used in the subsequent tracing experiments in the study.

### Neural tracing experiments

For initial surgery, following sedation (induction: ketamine (10mg Kg-^1^ IM)/midazolam (1mg Kg-^1^ IM), maintenance: isoflurane 1-2% inhalation) and intubation, a right unilateral thoracotomy was performed by dividing the pectoral muscle, making a small incision in the pericardium, and exposing the right atrial-superior vena cava junction. Then 5mg of Fast Blue (Polysciences), a neuronal tracer that is retained and diffuses within the lipid bilayer, in 250uL of sterile water (2% weight/volume) was injected using a 27-gauge needle into the SAN region. A chest tube was placed, and the incision was closed. Immediately prior to removal, the chest tube was aspirated. Tissues were harvested in a terminal procedure at least 3 weeks later as described below.

### Porcine tissue collection

Following sedation (induction: tiletamine-zolazepam 6mg Kg^−1^ IM, maintenance: isoflurane 1-2% inhalation) and intubation, we performed a midline sternotomy and exposed the heart. A heparin bolus of 5000U IV was administered, and the pig was then placed in ventricular fibrillation with application of a 9V battery to the surface of the heart. The heart was explanted and syringe-flushed with heparinized normal saline (5U mL^−1^) via the transected aorta. The area of interest (RAGP-SAN region) was then excised and rinsed in heparinized saline. RAGPs were separated from the SANs, immersed in 1x PBS at RT for 30 seconds and transferred to 25% Optimal Cutting Temperature compound (OCT, TissueTek; VWR 25608-930), followed by 50% OCT and then embedded in 100% OCT and placed in a cryomold. The cryomold was placed in a methanol dry ice bath for flash freezing.

### Cryosectioning and staining

RAGP was sectioned along the superior-inferior axis (corresponding to the source animal) at 40μm thickness, yielding between 447-1030 sections per RAGP, with corresponding blockface images. Tissue sections were mounted on PPS membrane slides (Leica Microsystems, Catalog 11600294). Slides were fixed in ice cold ethanol (100% Ethanol) for a minute followed by four minutes of staining with 0.0001% Cresyl Violet (ACROS Organics, AC229630050). The slides were dehydrated using 95% and 100% Ethanol followed by Xylene for one minute each. The staining protocol was kept to under 15 minutes and an RNAse inhibitor (Invitrogen SUPERase-In RNase Inhibitor, Catalog AM2696) was added to all aqueous reagents to preserve RNA quality. More detailed methods can be found in the research protocol available at DOI: 10.21203/rs.3.pex-928/v1.

### Laser capture microdissection

Slides were stained and immediately processed for sample collection to preserve RNA quality using Laser Capture Microdissection (Arcturus, ThermoFisher). SAN projecting and non-projecting neurons were identified under fluorescence using FastBlue (excitation 365 nm, emission 420 nm) and cresyl-violet (585 nm excitation and 627 nm emission))stain and only cresyl-violet stain respectively. Samples were collected on LCM Caps and stored at −80°C. Samples were lysed at the time of gene expression experiments with appropriate lysis reagents based on the downstream processing protocols (RNASeq or HT-qRTPCR). More detailed methods can be found in the research protocol available at DOI:10.21203/rs.3.pex-927/v1.

### Mapping LCM samples onto the reconstructed 3D stack

2D images of tissue blockface were acquired after each section. Using Tissue Mapper software 3D volume of each RAGP was created from the outer contours of blockface images. Images taken during LCM sample collection were assigned to corresponding sections in the 3D stack. Anatomical features and neurons (collected and not collected) were assigned markers enabling extraction of specific XYZ coordinates for the neurons in RAGP. Blockface images and acquisition images can be found at DOI: 10.26275/56h4-ypua. More detailed methods can be found in the research protocol available at DOI: 10.21203/rs.3.pex-922/v1.

### RNASeq library preparation

Samples on LCM caps were processed for single cell scale RNASeq based on a protocol modified from Foley et al., 2019^36^. We followed the steps recommended for fresh frozen tissue with the following modifications: We used 10μl of lysis buffer on HS Caps (original protocol recommends 5 μl). In the step 3 for performing PCR amplification of the tagged mRNA fragments, we used 22 cycles of PCR (maximum allowed). The original protocol relies on a formula to compute amplification cycles, and suggested 18 cycles based on sample RNA input and number of samples multiplexed per RNASeq run. Our initial tests determined this to be too low for yielding sufficient material for downstream sequencing. The remaining steps of the protocol were followed as in the original. More detailed methods can be found in the research protocol available at DOI: 10.21203/rs.3.pex-962/v1.

### Single cell RNA-seq data analysis

Raw sequencing data (Illumina sequencer’s base call files (BCLs) was converted to Fastq files using the Illumina bcl2fastq program. The reads were trimmed to remove poly A tail and G overhang, and the 5 base umi was extracted. The genome sequence was indexed and the single reads were aligned to the Sus scrofa reference genome sequence version Sus_scrofa.Sscrofa11.1.fasta available in the Ensembl database (RRID:SCR_002344), using STAR software (RRID:SCR_015899) STAR-2.7.2a^55^. A modified version of Sus_scrofa.Sscrofa11.1.95.gtf was used as a reference transcriptome. Feature count algorithm (featureCounts, RRID:SCR_012919), Subread R package^56^ was used to count the reads to genomic features - genes and exons. A digital gene expression matrix was created from the gene counts. Multiple batch correction algorithm ComBat-seq^57^ was used to account for technical variability arising from batch effect. Out of 142 samples, 52 Samples with non-zero gene counts <6000 were considered as outliers. Additionally, 10800 genes that are present in very low quantities (<30 non-zero gene counts) were filtered out. A regularized log transformation was carried out using DESeq2^58^. We normalized the filtered data using a quantile regression method SCnorm^59^. Our fInal matrix consisted of 90 samples and 15000 genes.

### Extraction of significant genes from GTEx

We generated a list of neuronal genes enriched in brain and neuronal tissues using data from the Genotype Tissue Expression (GTEx) project. We downloaded the median gene expression values for all tissue types from GTEx Analysis V8 (dbGaP Accession phs000424.v8.p2). Pavlidis Template Matching was used to find genes that were specifically enriched in brain and neuronal tissues (template-maximum for neuronal tissues, minimum for non-neuronal tissues) with p value < 0.01.

### Identification of highly variable genes using PCA

Principal component analysis (PCA) was performed on 90 neuronal RNAseq samples showing expression of 15,000 genes in the RAGP. 100 genes were taken from each of the first 50 PCs, the fifty genes most positively and most negatively contributing to each PC based on PC loadings, resulting in 1,814 genes that show the most variability across the RAGP.

### High-throughput real-time PCR

Single RAGP neurons in lysis buffer (Cells Direct Lysis Buffer, Invitrogen) were directly processed for reverse transcriptase reaction using SuperScript VILO Master Mix (Thermo Fisher Scientific, Waltham, MA), followed by real-time PCR for targeted amplification and detection using the Evagreen intercalated dye-based approach to detect the PCR-amplified product. Intron-spanning PCR primers were designed for every assay using Primer3^60^ and BLAST^61^. Genes were selected from across a wide array of neuronal functions, signal transduction and cell type identification. The standard BioMark protocol was used to process cDNA samples for 22 cycles of specific target amplification of 283 genes using TaqMan PreAmp Master Mix as per the manufacturer’s protocol (Applied Biosystems, Foster City, CA, USA). Real-time PCR reactions were performed using 96.96 BioMark Dynamic Arrays (Fluidigm, South San Francisco, CA, USA) enabling quantitative measurement of multiple mRNAs and samples under identical reaction conditions. Each run consisted of 30 amplification cycles (15 s at 95°C, 5 s at 70°C, 60 s at 60°C). Ct values were calculated by the Real-Time PCR Analysis Software (Fluidigm). Twenty one 96 × 96 BioMark Arrays were used to measure gene expression across the (422 samples before QC) single-cell samples. The same serial dilution sample set was included in each chip to verify reproducibility and test for technical variability. Samples from each animal were run across three chips to obtain data on 283 genes per sample. Each set of chip runs for a given animal contained overlapping assays that served as technical replicates to evaluate chip-to-chip variability. A chip-to-chip comparison of the serial dilution samples and neuronal/assay technical replicates demonstrates the high reproducibility with minimal technical variability of our data (Supplementary Fig. 7). More detailed methods can be found in the research protocol available at DOI: 10.21203/rs.3.pex-919/v1.

### HT-qPCR data analysis

Individual qRT-PCR results were examined to determine the quality of the qRT-PCR based on melt-curve analysis. Following this initial quality control, samples with >30% failed reactions and genes with >20% failed reactions were excluded from present analysis. A further 10 samples were determined to be outliers due overall gene expression distributions and were removed from the present analysis. Upon filtering based on these criteria, a total of 405 single-cell samples (152 non SAN-projecting neurons, 169 SAN-projecting neurons, and 84 neurons without reliable connectivity information) and 241 different gene assays were carried forward in the present analysis, with 211 genes showing >60% detectable expression across all RAGPs, which were used for the majority of the analysis. Raw Ct values for individual samples were normalized against a median expression level of a subset of 140 robustly expressed genes (genes with greater than 60% working reactions) across all animals to obtain -DCt values.The vector of median sample expression value was chosen over potential reference genes based on comparison of stable expression across all samples against known housekeeping genes using the ‘selectHKs’ function in the NormqPCR package in R (RRID:SCR_003388). The following equation was used to calculate -DCt values for each

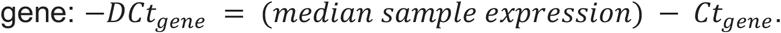

The -DCt data were then rescaled using the median across all samples within a gene using the following equation:

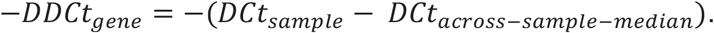

The raw and normalized dataset is available online as a Gene Expression Omnibus (RRID:SCR_005012) dataset (GEO reference ID: GSE149212) and on sparc.science (RRID:SCR_017041) portal at DOI: 10.26275/5jki-b4er.

### Determination of abundant/detectable expression

We considered a sample to have detectable expression of any given gene if any signal was detected in the HT-qPCR assay. A sample was considered to have abundant expression if its expression of that gene was less than 15 Ct. One exception is the expression level of Chat. The levels of Chat in the second female RAGP (PR1729) had a median expression of 7 Ct higher than that of the other 3 RAGP. Despite the reduced expression of Chat in raw Ct, it is important to note that the range of expression remains the same despite the decrease in abundance. Due to this consistency in range and pattern, we considered samples from PR1729 to have abundant expression if it had a Ct less than 22, consistent with the 7 Ct difference between the medians of expression.

### Clustering and template matching to identify transcriptional states

Single neurons were separated into different subpopulations based on their molecular profiles using Pavlidis Template Matching. Briefly, template matching estimates the correlation between a rescaled expression profile used as the template and a test expression profile. The canonical subpopulations were identified in SAN-projecting samples from one female and one male RAGP using hierarchical clustering (Pearson correlation, complete linkage) yielding seven sample clusters. The average silhouette width for all clusters (estimated using the cluster package in R) was 0.217, significantly higher than 1,000 randomized trials (Figure S3a). Gene medians of these clusters were used as templates to classify the non SAN-projecting cells into one of the canonical subpopulations. We used an R-based cutoff for our template match analysis (threshold R = 0.45). Single neurons that did not pass the R value threshold for any of the canonical templates were sequestered into a separate cluster, yielding 8 clusters total. Samples from another male (with cells annotated for projection) and female (without cells annotated for projection) were sorted into the original states based on correlation to the template profile (Figure 3a, S3b). Finally, based on visual examination, three states were combined to form state A based on their similar expression profiles, giving 6 total transcriptional states. An enrichment assessed by a one-way ANOVA and post hoc Tukey Honest Significant Difference was used to examine genes that were significantly enriched in one or more states. A gene was considered to be enriched in a state if p value < 0.01 for at least two other states (Figure 3d).

### Plots for showing abundance and range of gene expression in Ct space

In order to accurately display the range of the normalized data while also keeping in mind the overall abundance of each gene throughout the 4 RAGP, the −ΔΔCt data was visualized in Ct space. To do this, we first confirmed that the median Ct of each gene was relatively consistent across all 4 RAGP. The normalized expression for each animal was multiplied by −1 and then offset by the median gene expression across all 4 RAGP. This resulted in a plot that displayed expression level in Ct space (where the lower the Ct the higher the expression), showing the overall abundance of the gene in question across 4 RAGP while maintaining the normalized range of expression (across all RAGP for some figures, and comparing the ranges of each RAGP for other figures).

### Paracrine network analysis

Different paracrine signalling states of single neurons in the RAGP were created from the combination of identifiers that describe whether a neuron is more likely to transmit a neuropeptide signal, be activated by that signal, or both. Each neuron was identified as peptide+ or peptide-based on whether or not the neuron in question expressed the given peptide above or below the median expression value for that gene. Likewise, each neuron was identified as receptor+ if any one of the corresponding receptors showed expression above the median value for either receptor. Applying these identifiers to each peptide-receptor set resulted in 8 possible unique identifiers for each set, which when combined resulted in 64 possible unique identifiers. After applying these identifiers to the 405 sampled neurons, at least one neuron was assigned to 61 out of the 64 possible identifiers, indicating a large and widespread network of neuropeptide signalling, even when examining only 3 neuropeptides and their cognate receptors (Supplementary Cytoscape Network File). Of those 61 states, 12 groups have a frequency of more than 10 sampled neurons, representing a total of 224 out of the 405 single neurons, about 55%. A network representation of these 12 identified groups further highlights the interconnected nature of signalling between neurons.

### Confocal staining and visualization

Porcine RAGP fat pads were fixed in cold 4% paraformaldehyde in PBS for 24 h, cryoprotected in cold 20% sucrose in PBS, and sectioned at 30 mm thickness using a Leica CM3050S cryostat (Leica Microsystems Inc., Bannockburn, IL, USA). Sections were collected on charged slides and immunostained at room temperature using standard methods of fluorescence immunohistochemistry, as described previously^23,62,63^. Antibodies to vesicular acetylcholine transporter (VAChT; Synaptic Systems, Cat. No. 139103, RRID: AB_887864,1:500 dilution) and tyrosine hydroxylase (TH; Millipore, Cat. No. AB1542, RRID:AB_90755, 1:500 dilution) were used to label cholinergic and noradrenergic neurons, respectively. The VAChT antibody was generated in rabbit using aa 475-530 from rat VAChT as the immunogen, and it has been validated by experiments with knockout and knockdown mice as well as by immunohistochemistry and Western blotting. The TH antibody was generated in sheep using native TH from rat pheochromocytoma, and it has been validated by immunohistochemistry and Western blotting. Sections were washed and blocked before incubation overnight with both primary antibodies. This was followed by washing and blocking again before addition of species-specific donkey secondary antibodies conjugated to Alexa Fluor 488 for VAChT and biotin-SP for TH (Jackson ImmunoResearch Laboratories). After incubation for 2 h with secondary antibodies and washing with PBS, sections were incubated with streptavidin conjugated Cy3 (Jackson ImmunoResearch Laboratories) to amplify the TH signal. Sections were then washed with PBS, and cover glasses were applied with Citifluor (Ted Pella, Inc.) or SlowFade Gold antifade reagent (Life Technologies Corporation, Eugene, OR, USA). Cover glasses were sealed with clear nail polish. Specific staining did not occur in negative control sections processed without the addition of the primary antibodies.The localization of VAChT and TH was evaluated by confocal microscopy with a Leica TCS SP8 Confocal Microscope (Leica Microsystems Inc.). Confocal images were collected at a resolution of 1024X1024 by sequential scans using 488 and 552 laser lines. Stacks of optical sections, with the section number optimized by the software, were collected for regions of interest using 4-line averages for each scan. Stacks spanned the entire tissue thicknesses. Figures were created using maximum intensity projection images for individual channels and merged images. These images compress the stack of optical images into a two-dimensional view by showing only the highest intensity pixel across the stack for each point.

### Data Availability

The authors declare that all the data supporting the findings of this study are available within the article and its supplementary information files or from the corresponding author upon reasonable request. Raw sequencing data generated in this study have been deposited at the GEO database under accession code:GSE154119. Raw and processed HT-qPCR data of single neurons from the pig RAGP have been deposited in the GEO database under accession code: GSE149212. The RNAseq and HT-qPCR datasets constitute a GEO SuperSeries GSE154411. All sample acquisition images, raw and processed transcriptomic data, and annotations pertaining to 3D spatial location are publicly available in the sparc.science (RRID:SCR_017041) repository with the digital object identifiers 10.26275/56h4-ypua, 10.26275/kabb-mkvu, 10.26275/5jki-b4er, 10.26275/qkzi-b1mq, and 10.26275/255m-00nj. A dataset containing high-resolution figures, supplementary figures, movies and files, as well as the TissueMapper XML annotations and the R code to generate the data-driven plots and visualizations illustrated in the figures are available at 10.26275/jdws-d7md. Data analyzed from Hu et al. 2017 is available in the GEO database under the accession code GSE106678. Data from GTEX was retrieved from GTEx Analysis V8 (dbGaP Accession phs000424.v8.p2). Single cell data from the Human CNS was obtained through the Allen Brain Map (RRID:SCR_017001) project and is available at https://portal.brain-map.org/atlases-and-data/rnaseq/human-multiple-cortical-areas-smart-seq.

## Supporting information

Supplementary Cytoscape Networks

## Acknowledgements

Financial support for this work was provided by the National Institutes of Health under the Stimulating Peripheral Activity to Relieve Conditions (SPARC) program, Grant OT2 OD023848 (PI: K.S.) and NHLBI Grant U01HL133360 to J.S.S. and R.V. The authors thank Ms. Maci Heal for preparing the supplementary videos. The authors thank Dr. Anita Bandrowski for evaluating the manuscript through the SciScore™ tool to check for adherence to the Structure, Transparency, Accessibility and Reporting criteria and for supplying the RRIDs for relevant resources used in the study.

## Author Information

### Author Contributions

K.S., J.S., R.V. conceived, designed and supervised this work. S.R., S.A., S.T., and S.N. obtained and processed neuronal samples. P.H. and J.L.A. performed the neural tracing and tissue harvesting. A.M., S.A., and L.K. processed transcriptomic data for analysis. A.M. and S.A. analyzed HT-qPCR data. A.M. analyzed RNAseq data and external datasets. E.H.S. contributed to the acquisition, analysis and interpretation of immunohistochemical studies and drafting of the work. D.B.H. contributed to the conception of the work and analysis of immunohistochemical studies and drafting of the manuscript. A.M. and S.A. wrote and edited the manuscript with assistance from R.V. and J.S. A.M. and S.R. performed data visualizations and figure formulations. S.R. mapped all anatomy and neurons in 3D stacks.

### Ethics Declarations

Competing interests: University of California, Los Angeles has patents developed by KS and JLA relating to cardiac neural diagnostics and therapeutics. KS and JLA are cofounders of NeuCures, Inc. All the remaining authors declare no competing interests.

---

**Supplementary Figure 1:**
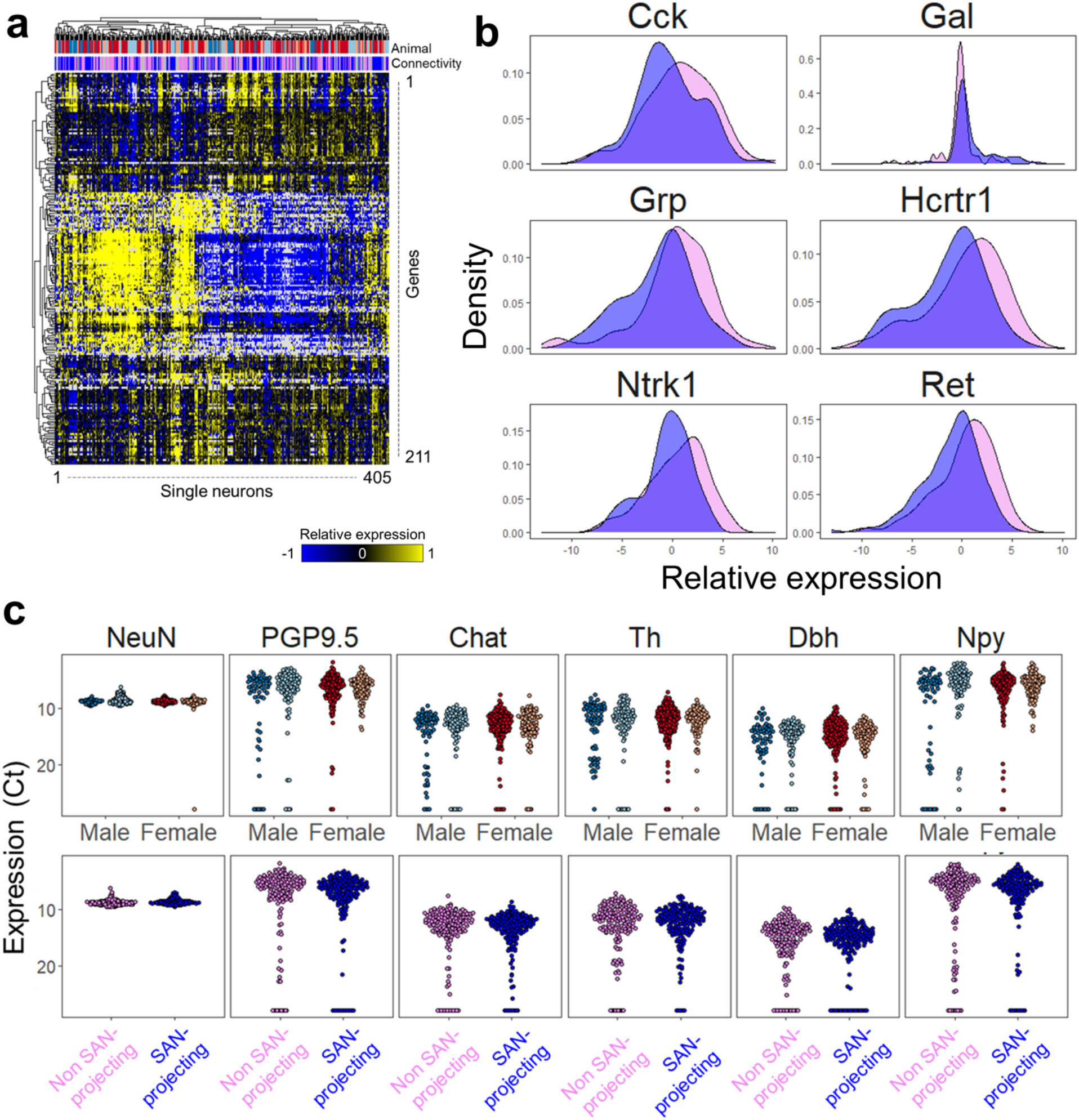
Expression of 405 single neuron samples across RAGP and between SAN-projecting and non SAN-projecting neurons. **(a)** Expression of 211 genes that have >60% detectable expression across 405 single neurons (n=4 animals). **(b)** Density distribution of SAN-projecting and non SAN-projecting single neurons for 6 genes that show statistical differences between projecting and non-projecting neurons (K-S statistic, FDR-adjusted *p* < 0.01, fold change > 2). **(c)** Expression distribution showing both the range and abundance of select neuronal markers across RAGP (top) and between SAN-projecting and non SAN-projecting neurons (bottom).

**Supplementary Figure 2:**
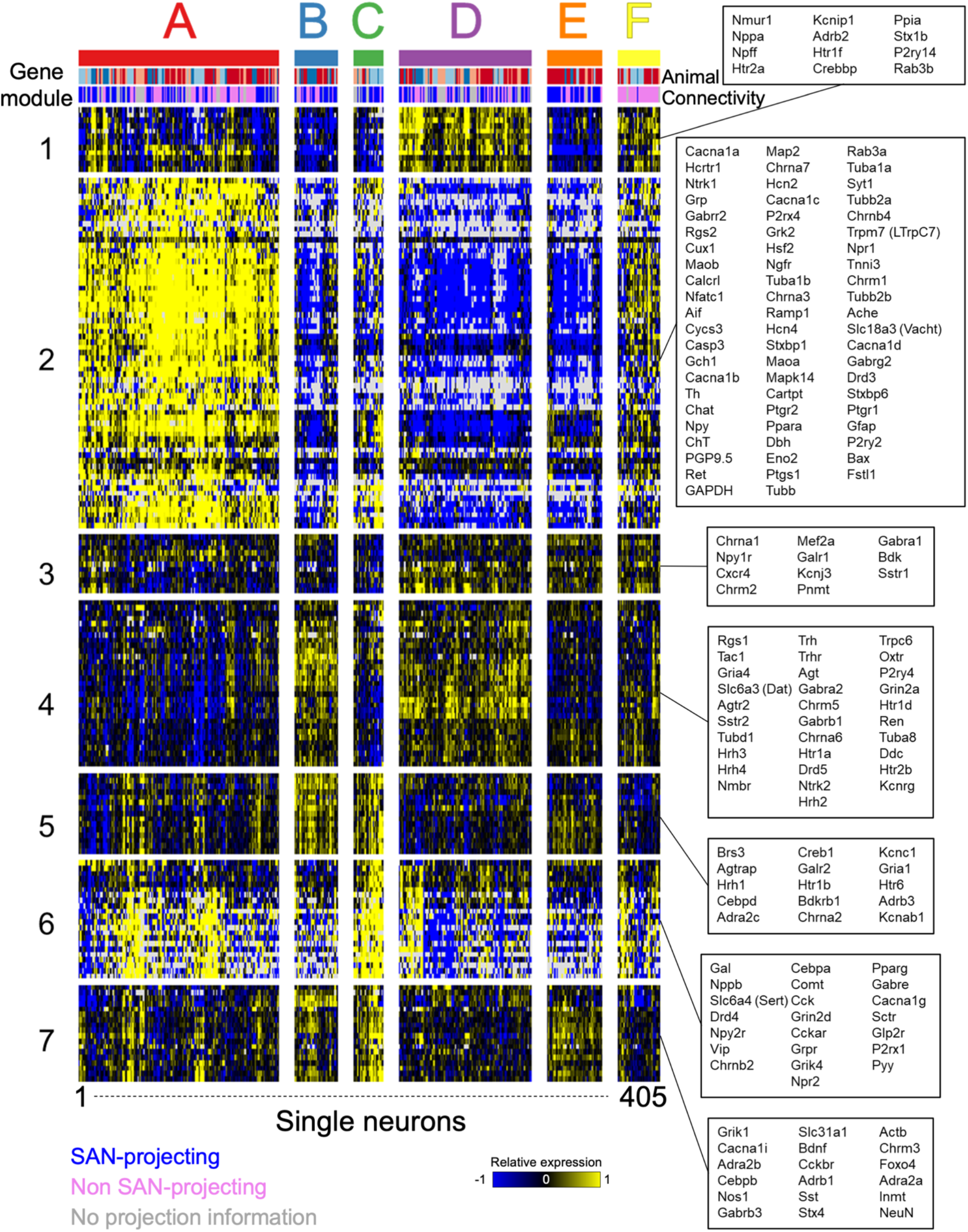
Landscape of neuronal transcriptional states across 4 RAGP. Expression of 174 genes each assayed in 405 single neurons (n=4 animals). Accompanies Fig. 4a which shows expression of 321 single neurons (n=3 animals). A fourth pig with no projection information was further incorporated into the identified transcriptional states. Genes names in each box are organized in the same order of the genes in the heatmap, to be read down each column.

**Supplementary Figure 3:**
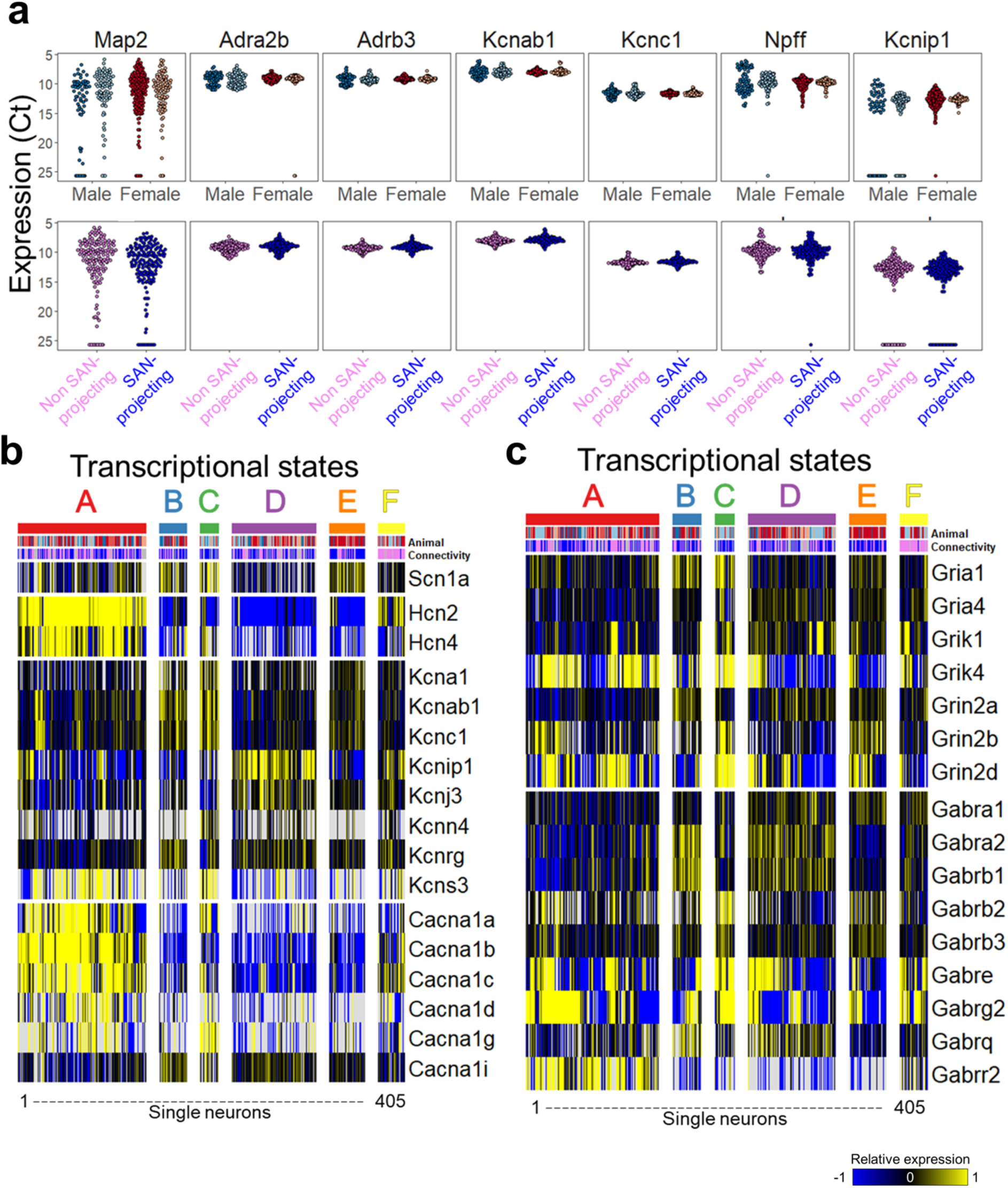
Expression patterns of key genes that differentiate transcriptional states and ion channels and transporters across single neurons in RAGP. **(a)** Expression distribution showing both the range and abundance for selected genes found to be enriched in a particular transcriptional state (Fig 4d-f). Expression is shown across RAGP (top) and between SAN-projecting and non SAN-projecting neurons (bottom). **(b-c)** Gene expression of various sodium, potassium, and calcium channels (b) and various glutamatergic and GABAergic receptors (c) across the identified transcriptional states.

**Supplementary Figure 4:**
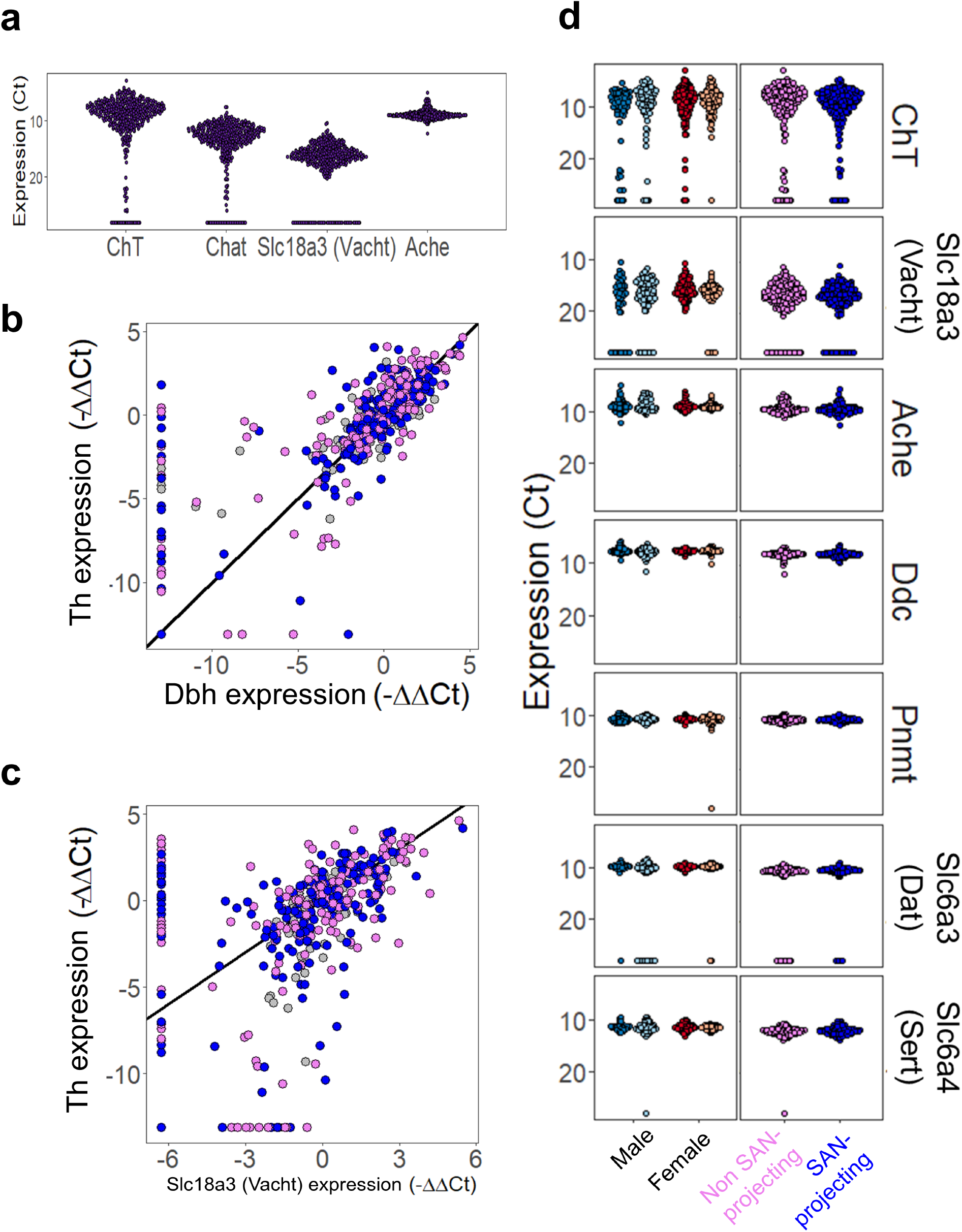
Expression of cholinergic and catecholaminergic markers. **(a)** Beeswarm plot showing the abundance and the range of expression of key genes involved in acetylcholine biosynthesis and transport across 405 single RAGP neurons (n=4 animals). **(b-c)** Co-expression of *Th* with *Dbh* (b) *and Slc18a3* (*Vacht*) (c) across 405 single RAGP neurons (n=4 animals). Solid line represents a diagonal of equal expression. **(d)** Expression distribution showing both the range and abundance for genes involved is biosynthesis and transport of catecholamines and acetylcholine. Expression is shown across RAGP (top) and between SAN-projecting and non SAN-projecting neurons (bottom). Remaining genes in these pathways can be found in Supplementary Fig 1a.

**Supplementary Figure 5:**
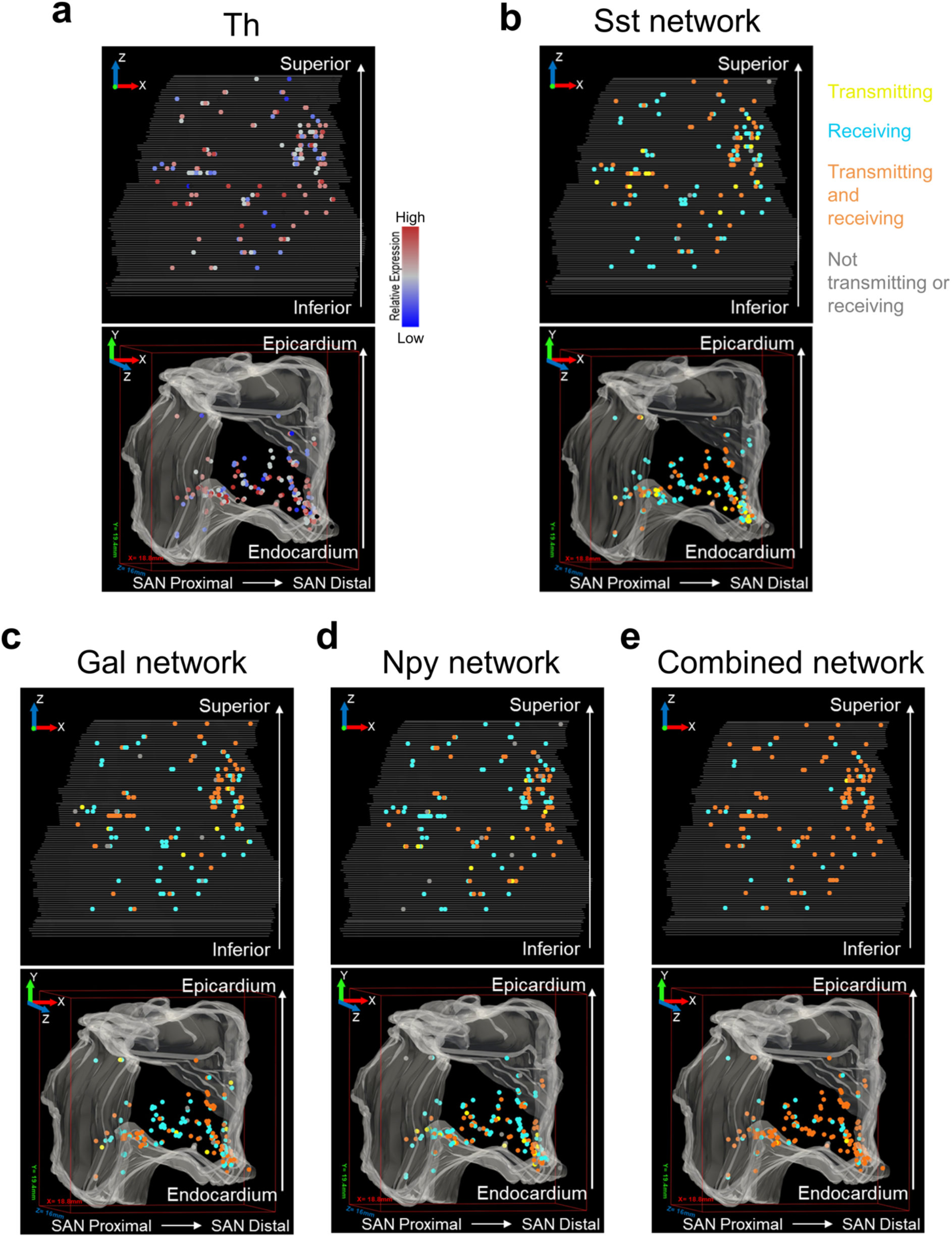
3D visualization of a representative RAGP. **(a)** Visualization of *Th* gene expression within the 3D anatomical framework for a representative RAGP. **(b-e)** Visualization of local paracrine networks for *Sst* (b), *Gal* (c), and *Npy* (d), as well as the combined network (e), within the 3D anatomical framework for a representative RAGP.

**Supplementary Figure 6:**
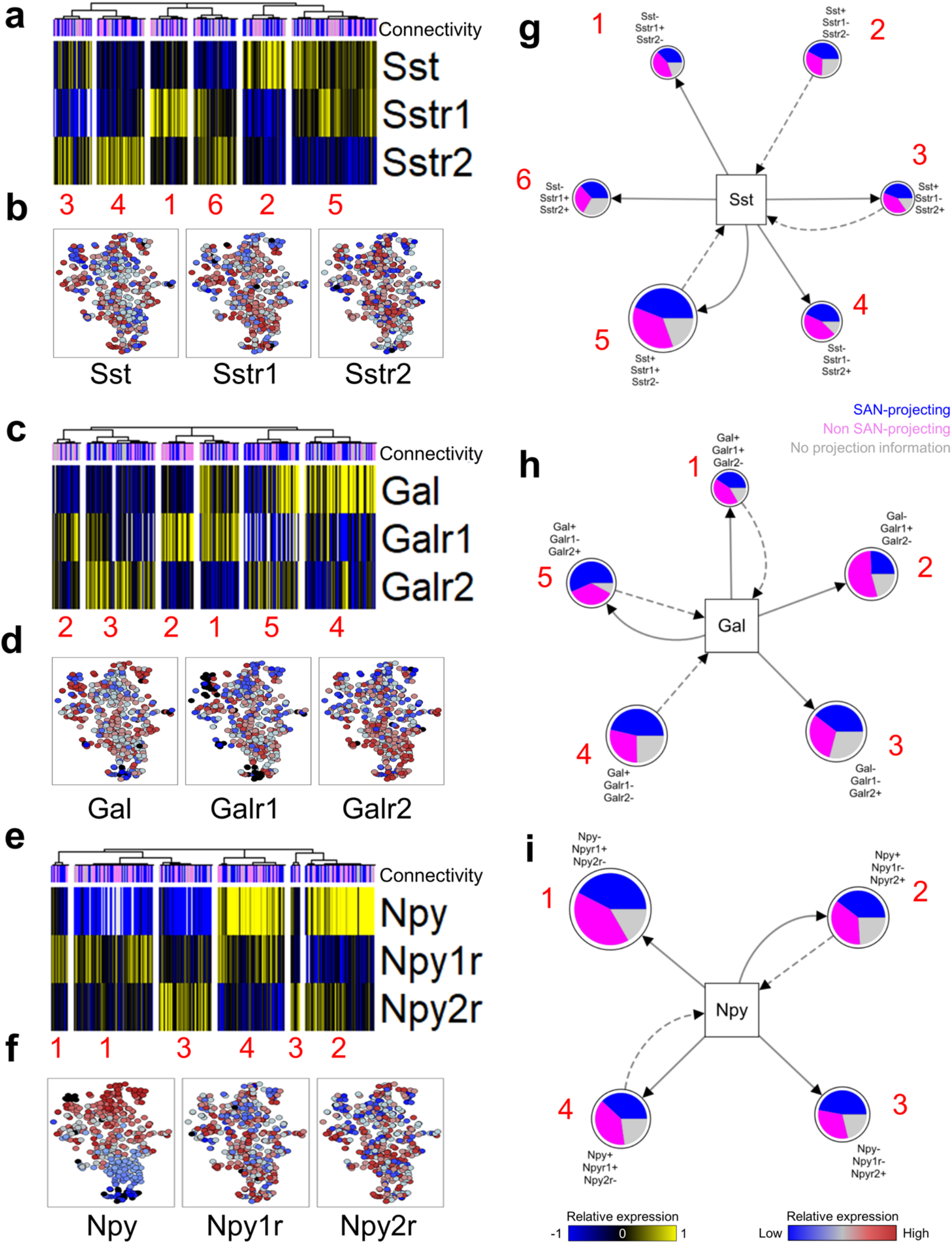
Neuropeptidergic interaction networks in the pig RAGP. **(a,c,e)** Expression patterns of somatostatin (a), galanin (c), and neuropeptide Y (e) and their cognate receptors across 405 single RAGP neurons (n=4 animals) that are numbered corresponding to their representative node in the networks for **(g-i). (b,d,f)** The transcriptional landscape across all RAGP colored for expression of the neuropeptides and their cognate receptors. **(g-i)** Network representations of the heatmaps from (a,c,e) where each cluster is represented as a specific node depending on whether or not it is transmitting the peptide signal or receiving it through one of the cognate receptors. The pie chart within each circular node indicates the proportion of the neurons within that subtype that are identified as projecting to the SAN region. The size of the node is proportional to the number of single neurons belonging to each subtype.

**Supplementary Figure 7:**
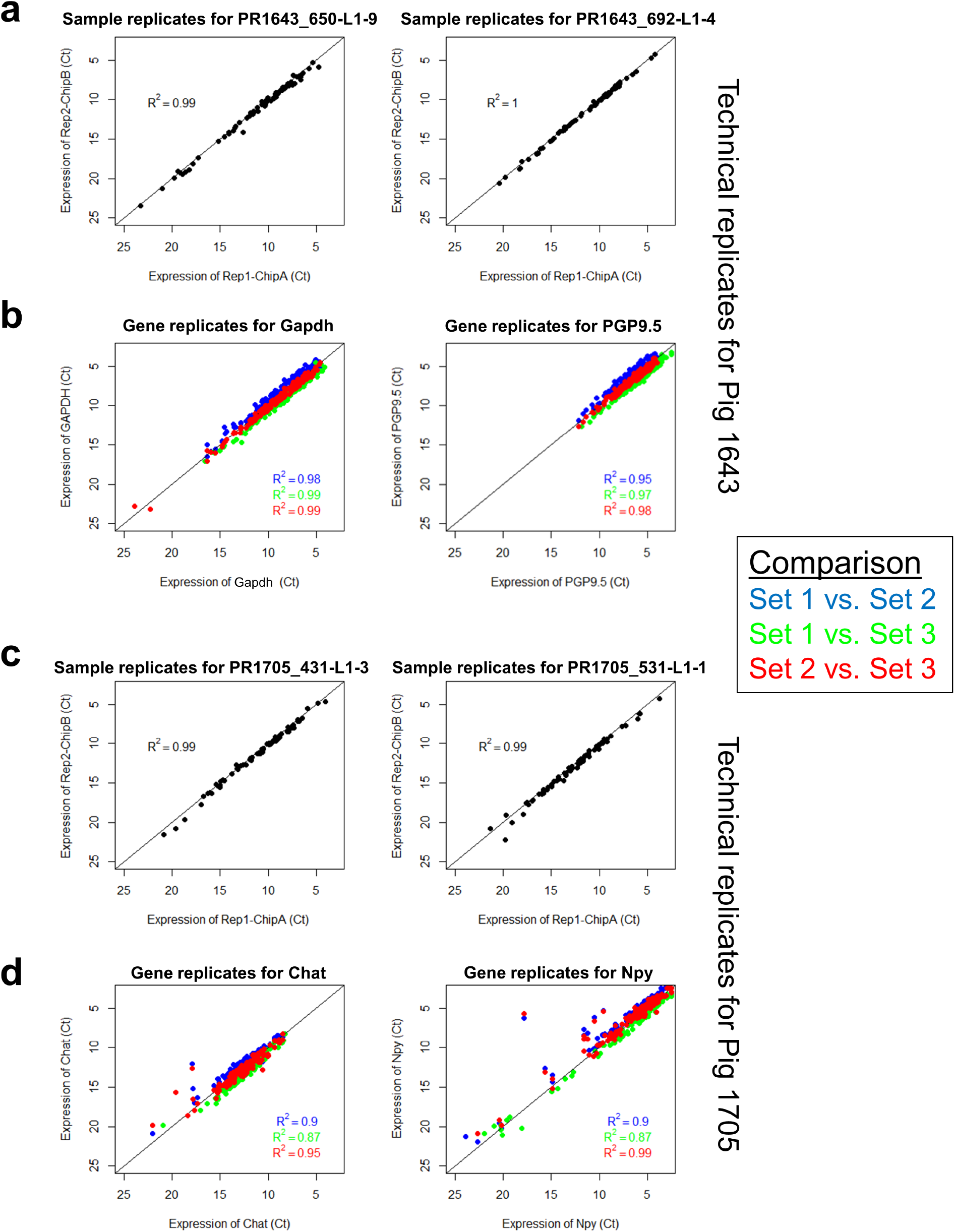
Technical reproducibility of HT-qPCR. **(a-d)** Technical Replicates of Samples and Genes in Pig 1643 (a-b) and Pig 1705 (c-d). To check for technical reproducibility within the HT-qPCR data, select samples and genes were run on multiple Biomark Chips. Samples had two replicates (a,c) and genes had three replicates (b,d). To robustly assess the three gene replicates, we compared each of the three sets to each other, shown in three different colors, where blue represents the comparison between sets 1 and 2, green represents the comparison between sets 1 and 3, and red represents the comparison between sets 2 and 3.

**Supplementary Movie 1:**

Acquisition of single neurons from the RAGP and 3D visualization of SAN-projecting and non-SAN-projecting neurons within a representative RAGP.

**Supplementary Movie 2:**

Visualization of identified transcriptional states within the 3D anatomical framework of a representative RAGP.

**Supplementary Movie 3:**

3D spatial distribution of *Chat* and *Th* expression across a representative RAGP.

**Supplementary Movie 4:**

Neuropeptidergic interaction networks within the 3D anatomical framework of a representative RAGP.

**Supplementary Cytoscape Network File:**

Cytoscape file containing all the network information related to Figure 6 and Supplementary Movie 4. This Cytoscape session file includes annotated networks corresponding to the individual neuropeptides as well as the combined neuropeptidergic network mentioned in the text and the simplified network shown in Figure 6.

